# Quantifying the potential for red blood cell β-adrenergic sodium-proton exchangers to protect oxygen transport in hypoxic and hypercapnic white seabass

**DOI:** 10.1101/2021.04.28.441819

**Authors:** Till S. Harter, Alexander M. Clifford, Martin Tresguerres

## Abstract

White seabass (*Atractoscion nobilis*) are increasingly experiencing periods of low oxygen (O_2_; hypoxia) and high carbon dioxide (CO_2_, hypercapnia) due to climate change and eutrophication of the coastal waters of California. Hemoglobin (Hb) is the principal O_2_ carrier in the blood and in many teleost fishes Hb-O_2_ binding is compromised at low pH; however, the red blood cells (RBC) of some species regulate intracellular pH with adrenergically-stimulated sodium-proton-exchangers (β-NHE). We hypothesized that RBC β-NHEs in white seabass are an important mechanism that can protect the blood O_2_-carrying capacity during hypoxia and hypercapnia. We determined the O_2_-binding characteristics of white seabass blood, the response of RBCs to adrenergic stimulation, and quantified the protective effect of β-NHE activity on Hb-O_2_ saturation. White seabass had typical teleost Hb characteristics, with a moderate O_2_ affinity (PO_2_ at half-saturation; P_50_ 2.9 kPa) that was highly pH-sensitive (Bohr coefficient -0.92; Root effect 52%). The presence of RBC β-NHEs was confirmed by functional, molecular and bioinformatic data and super-resolution imaging revealed, for the first time, the subcellular location of β-NHE protein in vesicle-like structures and on the RBC membrane, and its translocation after adrenergic stimulation. The activation of RBC β-NHEs increased Hb-O_2_ saturation by ∼8% in normoxia at 1 kPa PCO_2_, and by up to 20% in hypoxia. Our results confirm that RBC β-NHE activity in white seabass has great potential to protect arterial O_2_ transport in environmentally relevant conditions of hypoxia and hypercapnia, but also reveal a potential vulnerability of fish to combinations of these stressors.

## Introduction

White seabass (*Atractoscion nobilis*), a teleost fish species endemic to the coastal waters of California, are apex-predators with ecological significance, sought-after targets of recreational and commercial fisheries and are gaining importance in aquaculture. Their natural habitat along the kelp forests of the north-eastern Pacific is subject to strong seasonal fluctuations in water chemistry, due to the upwelling of deeper waters that are often depleted of oxygen (O_2_; hypoxia), have a high carbon dioxide tension (CO_2_; hypercapnia) and thus, a low pH (1). In addition, a steadily warming climate and growing anthropogenic nutrient loading are increasing the frequency of large algae blooms (“red tides”) in California’s coastal waters (2). When these algae blooms wane, the microbial decomposition of their biomass, consumes O_2_ and produces CO_2_ and other toxic and acidic by-products of biological decay, such as hydrogen sulfide, altogether creating large hypoxic and hypercapnic zones (3). Species that are highly mobile may be able to avoid these areas, but for many sedentary species, survival will depend on enduring these conditions. Climate change at large is also leading to more hypoxic and acidic oceans and, over generations, some species may adapt to cope with their altered habitats (4). However, the reoccurrence of upwelling and severe algae blooms may acutely expose animals to conditions that far exceed worst-case predictions for the end of the century, creating strong selective pressures for hypoxia and hypercapnia tolerance and perhaps overwhelming the rates at which some species can adapt to climate change.

The most recent severe red tide in Southern California occurred in April-May of 2020, when the water measurements at the pier of the Scripps Institution of Oceanography (SIO) in La Jolla (CA, USA) revealed average daily dissolved O_2_ levels of <2 mg l^-1^ and pH as low as 7.06 (5). At a water temperature of 17°C and 35 ppt salinity, these values correspond to 5.3 kPa PO_2_ (6) and 1.16 kPa PCO_2_ (CO2SYS software;, 7). Equally alarming was the prolonged duration of hypoxia, where for nine consecutive days water PO_2_ was below the threshold (4.6 mg l^-1^) that is considered lethal for 90% of marine life (8). The SIO aquatics facility is supplied with water taken in at the pier, which resulted in hypoxic and hypercapnic exposures of all research animals, despite every effort to aerate the tanks. However unfortunate, this natural experiment revealed a remarkable tolerance of white seabass to these adverse water conditions, despite being deprived the behavioral avoidance of hypoxia that may be recruited in the wild. Therefore, the aim of the present study was to explore the O_2_-transport capacity of white seabass with a focus on the cellular mechanisms at the level of the red blood cell (RBC) that may contribute to their hypoxia and hypercapnia tolerance.

For obligate aerobic animals, the challenge to surviving unavoidable environmental hypoxia is balancing the uptake and delivery of O_2_ with its consumption in the mitochondria (9). Hemoglobin (Hb) is the principal O_2_ carrier in the blood and therefore the cardiovascular O_2_-carrying capacity is largely determined by the O_2_-binding characteristics of Hb. As such, a higher Hb-O_2_ affinity will favor the extraction of O_2_ from hypoxic waters and thus, a lower Hb P_50_ (the partial pressure of O_2_ at which Hb is 50% saturated) is typically associated with hypoxia tolerance in fishes (10, 11); however, whether white seabass have high-affinity Hbs that would confer some hypoxia tolerance is currently unknown.

Hb-O_2_ binding in teleost fishes is highly pH-sensitive, where a reduction in pH decreases Hb-O_2_ affinity via the Bohr effect (12), and the Root effect prevents Hb from becoming fully O_2_-saturated at low pH, even at super-atmospheric PO_2_ (13, 14). The reduction in Hb-O_2_ carrying capacity due to the Root effect is physiologically significant, as it enhances the unloading of O_2_ at the eyes and the swimbladder of teleosts, where blood is acidified locally (15–17). In contrast, during a systemic blood acidosis that may occur during exercise or hypoxia, the pH-sensitive Hbs of teleosts may fail to become fully oxygenated at the gills, decreasing the O_2_-carrying capacity of arterial blood and leading to hypoxemia at the tissues. Thus, a combined hypoxic and hypercapnic exposure may be especially dangerous for teleosts, as a reduced availability of O_2_ in the environment is paired with the simultaneous reduction of Hb-O_2_ affinity via the Bohr effect at low pH; however, whether white seabass have pH-sensitive Hbs is currently unknown.

Hb is housed within RBCs that, in teleosts, may prevent systemic hypoxemia by actively regulating their intracellular pH (pH_i_) to protect Hb-O_2_ binding during a reduction in extracellular pH (pH_e_). In brief, a decrease in arterial PO_2_ or pH leads to the release of catecholamines into the blood (18, 19), which bind to a β-adrenergic receptor on the RBC membrane and activate a sodium-proton-exchanger (β-NHE, Slc9a1b,) via the cyclic adenosine monophosphate (cAMP) pathway (20). The extrusion of H^+^ by the β-NHE raises pH_i_ above the equilibrium condition, which increases Hb-O_2_ affinity and will promote the extraction of O_2_ from hypoxic waters (21). The adrenergic stimulation of RBCs also causes an influx of Na^+^ and Cl^-^ that leads to osmotic swelling and that has been used as a marker to determine the presence of RBC β-NHEs in fish species (22, 23). A broader phylogenetic analysis indicates that most teleosts, but not other fishes, have RBC β-NHEs (24); however, whether white seabass RBCs have β-NHE activity is currently unknown.

Based on these considerations, we hypothesized that β-NHE activity in white seabass is an important mechanism that can protect the blood O_2_-carrying capacity during environmentally relevant levels of hypoxia and hypercapnia (PO_2_<5.3 kPa and PCO_2_ <1.16 kPa; see above). We tested this hypothesis in a series of *in vitro* experiments and predicted that: i) white seabass have a high Hb-O_2_ affinity to maintain O_2_ uptake under hypoxic conditions, which was addressed by generating oxygen equilibrium curves (OEC) over a range of PO_2_; ii) white seabass display the large Bohr and Root effects that are typical of teleosts, which was addressed by generating OECs over a range of PCO_2_, and measuring pH_e_ and RBC pH_i_; iii) white seabass have a RBC β-NHE, which was addressed using molecular, bioinformatic and immunocytochemical techniques to establish its presence and localization, and by measuring RBC swelling after adrenergic stimulation and the inhibition of NHEs with amiloride; and finally iv) we quantified the protective effect of RBC β-NHE activity on blood O_2_-carrying capacity under environmentally relevant conditions, by measuring Hb-O_2_ saturation at increasing levels a hypercapnia in normoxia and hypoxia.

## Materials and Methods

### Animals and husbandry

White seabass (*A. nobilis*, Ayres 1860) were obtained from the Hubbs Sea World Research Institute (HSWRI, Carlsbad, USA) and were held indoors at the SIO aquatics facility for several months before experiments. Photoperiod was set to a 12:12 h light-dark cycle and fish were housed in large fiberglass tanks (∼3.5-10 m^3^) supplied with flow-through seawater from an inshore intake; the average water temperature at the time of experiments was 17°C. Aeration was provided to ensure normoxic conditions in all tanks (>90% air saturation of O_2_) and these water parameters were monitored every day. All fish were fed twice a week with commercial dry pellets (Skretting; Classic Bass 9.5 mm; Stavanger, Norway) and feeding was suspended 48 h before blood sampling. The white seabass used for the determination of blood O_2_-binding characteristics had an average weight of 1146±96 g (*N* = 8), while those used for the β-NHE experiments had an average weight of 357±27 g (*N* = 6). Animal husbandry and all experimental procedures were in strict compliance with the guidelines by the Institutional Animal Care and Use Committee (IACUC) and approved by the Animal Care Program at the University of California San Diego (Protocol no. S10320).

### Blood sampling

White seabass were moved individually into darkened boxes supplied with air and flow-through seawater, 24 h prior to blood sampling. The next day the water supply was shut off and the fish were anesthetized by carefully pouring a diluted benzocaine solution (Fisher Scientific, Acros 150785000; Waltham, USA; concentrated stock made up in ethanol) into the box without disturbing the fish, for a final concentrations of 70 mg l^-1^ benzocaine (<0.001% ethanol). After visible loss of equilibrium, fish were transferred to a surgery table, positioned ventral-side-up and their gills were perfused with water containing a maintenance dose of anesthetic (30 mg l^-1^ benzocaine). Blood sampling was by caudal puncture and 3 ml of blood were collected into a heparinized syringe. This procedure ensured minimal disturbance of the fish (25), which can decrease blood pH due to air-exposure (respiratory acidosis) and due to anaerobic muscle contractions during struggling (metabolic acidosis). After sampling, the fish were recovered and returned to their holding tank, and each individual was only sampled once. In the lab, the blood was centrifuged at 500 *g* for 3 min to separate the plasma from the blood cells. The plasma was collected in a bullet tube and stored over-night at 4°C. To remove any catecholamines released during sampling, the blood cells were rinsed three times in cold Cortland’s saline (in mM: NaCl 147, KCl 5.1, CaCl 1.6, MgSO_4_ 0.9, NaHCO_3_ 11.9, NaH_2_PO_4_ 3, glucose 5.6; adjusted to the measured plasma characteristics in white seabass of 345 mOsm and pH 7.8) and the buffy coat was aspirated generously to remove white blood cells and platelets. Finally, the pellet was re-suspended in 10 volumes of fresh saline to allow the RBCs to return to a resting state and stored aerobically on a tilt-shaker, over-night, at 17°C (26).

### Blood O_2_-binding characteristics

The next day, RBCs were rinsed with saline three times and re-suspended in their native plasma at a hematocrit of 5%; this value was chosen based on preliminary trials and yields an optic density that allows for spectrophotometric measurements of Hb-O_2_ binding characteristics (∼0.6 mM Hb). A volume of 1.4 ml of blood was loaded into a glass tonometer at 17°C and equilibrated to arterial gas tensions (21 kPa PO_2_, 0.3 kPa PCO_2_ in N_2_) from a custom-mixed gas cylinder (Praxair; Danbury, USA). After one hour, 2 µl of blood were removed from the tonometer and loaded into the diffusion chamber of a spectrophotometric blood analyzer (BOBS, Loligo Systems; Viborg, Denmark). The samples were equilibrated to increasing PO_2_ tensions (0.5, 1, 2, 4, 8, 16 and 21 kPa PO_2_) from a gas mixing system (GMS, Loligo), in two-minute equilibration steps and the absorbance was recorded once every second at 190-885 nm. At the beginning and end of each run, the sample was equilibrated to high (99.7 kPa PO_2_, 0.3 kPa PCO_2_ in N_2_ for 8 min) and low (0 kPa PO_2_, 0.3 kPa PCO_2_ in N_2_ for 8 min) PO_2_ conditions; for the calculation of Hb-O_2_ saturation from raw absorbance values, it was assumed that Hb was fully oxygenated or deoxygenated under the two conditions, which was confirmed by inspecting the absorption spectra (all raw data are deposited online). A PCO_2_ of 0.3 kPa was maintained throughout these trials to prevent RBC pH_i_ from increasing above physiologically relevant levels and this value was chosen to match that measured in the arterial blood of rainbow trout (*Oncorhynchus mykiss*) *in vivo* (27). All custom gas mixtures were validated by measuring PO_2_ with an FC-2 Oxzilla and PCO_2_ with a CA-10 CO_2_ analyzer (Sable Systems, North Las Vegas, USA) that were calibrated daily against high purity N_2_, air, or 5% CO_2_ in air.

An additional 250 µl of blood were removed from the tonometer to measure blood parameters as follows. Hematocrit (Hct) was measured in triplicate in microcapillary tubes (Drummond Microcaps, 15 µl; Parkway, USA), after centrifuging at 10,000 *g* for 3 min. Hb was measured in triplicate using the cyano-methemoglobin method (Sigma-Aldrich Drabkin’s D5941; St. Louis, USA) and an extinction coefficient of 10.99 mmol cm^-1^ (28). Blood pH was measured with a thermostatted microcapillary electrode at 17°C (Fisher Accumet 13-620-850; Hampton, USA; with Denver Instruments UB-10 meter; Bohemia, USA), calibrated daily against precision pH buffers (Radiometer S11M007, S1M004 and S11M002; Copenhagen, Denmark). Thereafter, the blood was centrifuged to separate plasma and RBCs and total CO_2_ content (TCO_2_) of the plasma was measured in triplicate with a Corning 965 (Midland, USA). The RBCs in the pellet were lysed by three freeze-thaw cycles in liquid nitrogen and pH_i_ was measured in the lysate as described for pH_e_ (29). After completing these measurements, the PCO_2_ in the tonometer was increased in steps from 0.3 to 2.5 kPa and, each time, OECs and blood parameters were measured as described above.

### RBC swelling after β-adrenergic stimulation

After storage of blood samples over-night in saline, the RBCs were rinsed three times in fresh saline and re-suspended in their native plasma at a Hct of 25%. A volume of 1.8 ml was loaded into a tonometer and equilibrated to 3 kPa PO_2_ and 1 kPa PCO_2_ in N_2_ at 17°C for one hour; similar hypoxic and acidotic conditions have been shown to promote β-NHE activity in other teleost species (30, 31). After one hour, an initial subsample of blood was taken and Hct, Hb, pH_e_ and pH_i_ were measured as described above. Thereafter, the blood was split into aliquots of 600 µl that were loaded into individual tonometers and treated with either: i) a carrier control (0.25% dimethyl sulfoxide, DMSO; VWR BDH 1115; Radnor, USA), ii) the β-adrenergic agonist isoproterenol (ISO; Sigma I6504; 10 µM final concentration, which stimulates maximal β-NHE activity in rainbow trout;, 19), or iii) ISO plus the NHE inhibitor amiloride (ISO+Am; Sigma A7410; 1 mM, according to 32). These treatments were staggered so that samples from each tonometer could be taken for the measurements of blood parameters at 10, 30 and 60 mins after drug additions.

To collect RBC samples for immunocytochemistry, the above tonometry trial was repeated with RBCs that were suspended in saline instead of plasma; this step was necessary as initial trials showed that plasma proteins interfered with the quality of cell fixations. Subsamples were removed from individual tonometers at the initial and 60 min time points. A volume of 60 µl was immediately re-suspended in 1.5 ml ice-cold fixative (3% paraformaldehyde, 0.175 % glutaraldehyde in 0.6 x phosphate buffered saline with 0.05 M sodium cacodylate buffer; made up from Electron Microscopy Sciences RT15949, Hatfield, USA) and incubated for 60 min on a revolver rotator at 4°C. After fixation, cells were washed three times in 1 x phosphate Buffered Saline (PBS, Corning 46-013-CM, Corning, USA) and stored at 4°C for processing (after visual inspection of cell morphology the fixation resulted in satisfactory results for *N* = 4 out of 6 fish). An additional subsample of 100 µl was removed from the tonometers and centrifuged to remove the saline. The RBC pellet was re-suspended in 5 volumes of lysis buffer containing 1 mM DL-Dithiothreitol (DTT; Thermo Fisher R0861; Waltham, USA), 1 mM phenylmethylsulfonyl fluoride (PMSF; Sigma P7626) and 10 mM benzamidine hydrochloride hydrate (BHH; Sigma B6506) in PBS. The RBCs were lysed by three cycles of freeze-thawing in liquid nitrogen, the lysate was centrifuged at 500 *g* for 10 min at 4°C and the supernatant was frozen at -80°C for Western blot processing.

### Hb-O_2_ binding after β-adrenergic stimulation

An additional aliquot of RBCs was re-suspended in native plasma at a Hct of 5%. Volumes of 300 µl were loaded into one of four tonometers and equilibrated to arterial gas tensions at 17°C (as described previously) and treated with either: i) a carrier control (DMSO; 0.25%), ii) ISO (10 µM), or iii) ISO+Am (1 mM). These treatments were staggered to allow for standardized measurements at 60 min after drug additions, when 2 µl of blood were removed from the tonometer and loaded into the BOBS for real-time measurements of Hb-O_2_ saturation during a respiratory acidosis. Therefore, the blood was exposed to stepwise increases in PCO_2_ (0.3, 0.5, 1, 1.5, 2 and 3 kPa) allowing for two minutes of equilibration at each step; preliminary trials showed full equilibration to the new PCO_2_ after ∼1 min and absorbance remained constant thereafter. As described previously, this protocol also included initial and final calibration steps, at which the sample was fully O_2_-saturated and then desaturated. A first trial was performed in normoxia (21 kPa PO_2_) and then a second sample was loaded form the same tonometer for an additional run under hypoxic conditions (3 kPa PO_2_). The PO_2_ value in these hypoxic runs was chosen to yield a Hb-O_2_ saturation close to P_50_ and was informed from the previous measurements of Hb-O_2_ binding characteristics. Finally, 250 µl of blood were removed from the tonometer for the measurement of blood parameters, as described previously.

### Subcellular localization of RBC β-NHE

Fixed RBCs were permeabilized in 1.5 ml 0.1% triton-X100 (VWR Amresco 1421C243) in PBS for 15 min at room temperature on a revolver rotator. Thereafter, the RBCs were blocked for auto-fluorescence in 100 mM glycine in PBS for 15 min, after which the cells were rinsed three times in PBS. For immunocytochemistry, 200 µl of these fixed RBCs were re-suspended in a blocking buffer containing 3% bovine serum albumin (VWR 0332) and 1% normal goat serum (Lampire Biological Laboratories 7332500; Pipersville, USA) in PBS and incubated for six hours on a rotator. Primary antibodies were added directly into the blocking buffer and incubated on a rotator over-night at 4°C. A monoclonal mouse anti-*Tetrahymena* α-tubulin antibody (deposited by Frankel, J. and Nelsen, E.M at the Developmental Studies Hybridoma Bank, DSHB12G10; Iowa City, USA) was used at 0.24 µg ml^-1^ and a custom, polyclonal, affinity-purified, rabbit anti-rainbow trout β-NHE (epitope: MERRVSVMERRMSH) was used at 0.02 µg ml^-1^. After primary incubations the RBCs were washed three times in PBS and incubated for three hours on a rotator at room temperature in blocking buffer containing secondary antibodies: 1:500 goat anti-mouse (AlexaFlour 568; Thermo Fisher Life Technologies A-11031), 1:500 goat anti-rabbit (AlexaFlour 488; A-11008) and 1:1000 4′,6-diamidino-2-phenylindole (DAPI; Roche 10236276001; Basel, Switzerland). After secondary incubations, RBCs were washed three times and were re-suspended in PBS. To validate the β-NHE antibody, controls were performed by leaving out the primary antibody and by pre-absorbing the primary antibody with its pre-immune-peptide. All images were acquired with a confocal laser-scanning fluorescence microscope (Zeiss Observer Z1 with LSM 800, Oberkochen, Germany) and ZEN blue edition software v.2.6. For super-resolution imaging the cells were re-suspended in PBS with a mounting medium (Thermo Fisher Invitrogen ProLong P36980) and acquisition was with the Zeiss AiryScan detector system. To ensure that images were comparable, the acquisition settings were kept identical between the different treatments and between treatments and the controls. Optical sectioning and three-dimensional (3D) reconstructions of single RBCs from the different treatments were processed with the Imaris software v.9.0. (Oxford Instruments, Abingdon, UK) and rendered into movies.

### Molecular β-NHE characterization

For Western blotting, RBC crude homogenates were thawed and centrifuged at 16,000 *g* for one hour at 4°C to obtain a supernatant containing the cytoplasmic fraction and a pellet containing a membrane-enriched fraction that was re-suspended in 100 µl of lysis buffer. The protein concentration of all three fractions was measured with the Bradford’s assay (BioRad 5000006; Hercules, USA). Samples were mixed 1:1 with Laemmli’s sample buffer (BioRad 161-0737) containing 10% 2-Mercaptoethanol (Sigma M3148) and were heated to 75°C for 15 min. Sample loading was at 5 µg protein from each fraction for the detection of β-NHE and 60 µg protein from crude homogenate for the detection of α-tubulin, into the lanes of a 5% stacking-and 10% separating polyacrylamide gel (Biorad, MiniProtean Tetra cell). The proteins were separated at 60 V for 30 min and 150 V until the Hb fraction (∼16 kDa) ran out the bottom of the gel (∼60 min); previous trials had shown that the high Hb content of these lysates may bind some antibodies non-specifically. The proteins were transferred onto a Immun-Blot polyvinylidene difluoride membrane (PVDF; BioRad) using a semi-dry transfer cell (Bio-Rad Trans-Blot SD) over-night, at 90 mA and 4°C. PVDF membranes were blocked over-night, on a shaker at 4°C in tris-buffered saline with 1% tween 20 (TBS-T; VWR Amresco ProPure M147) and 0.1 g ml^-1^ skim milk powder (Kroger; Cincinnati, USA). Primary antibodies were made up in blocking buffer and mixed on a shaker at 4°C, over-night, before applying to the membranes. The anti-α tubulin antibody was used at 4.7 ng ml^-1^, the anti-β-NHE antibody at 0.42 ng ml^-1^, and controls at a peptide concentration exceeding that of primary antibody by 10:1. Primary incubations were for four hours on a shaker at room temperature and membranes were rinsed three times in TBS-T for 5 min. Secondary incubations were with either an anti-rabbit or mouse, horse-radish peroxidase conjugated secondary antibody (BioRad 1706515 and 1706516) for three hours on a shaker at room temperature. Finally, the membranes were rinsed three times in TBS-T for 5 min and the proteins were visualized by enhanced chemiluminescense (BioRad, Clarity 1705061) in a BioRad Universal III Hood with Image Lab software v.6.0.1. Protein sizes were determined relative to a precision dual-colour protein ladder (BioRad 1610374).

The white seabass β-NHE sequence was obtained by transcriptomics analysis of gill samples that were not perfused to remove the blood and these combined gill and RBC tissue samples were stored in RNA later for processing. Approximately 50 µg of sample were transferred into 1 ml of Trizol reagent (Thermo Fisher 15596026) and were homogenized on ice with a handheld motorized mortar and pestle (Kimble Kontes, Dusseldorf, Germany). These crude homogenates were centrifuged at 1000 *g* for 1 min and the supernatant was collected for further processing. RNA was extracted in RNA spin columns (RNAEasy Mini; Qiagen, Hilden, Germany) and treated with DNAse I (ezDNase; Thermo Fisher, 11766051) to remove traces of genomic DNA. RNA quantity was determined by spectrophotometry (Nanodrop 2000; Thermo Fisher) and RNA integrity was determined with an Agilent 2100 Bioanalyzer (Agilent; Santa Clara, USA). Poly-A enriched complementary DNA (cDNA) libraries were constructed using the TruSeq RNA Sample Preparation Kit (Illumina; San Diego, USA). Briefly, mRNA was selected against total RNA using Oligo(dt) magnetic beads and the retained RNA was chemically sheared into short fragments in a fragmentation buffer, followed by first- and second-stand cDNA synthesis using random hexamer primers. Illumina adaptor primers (Forward P5-Adaptor, 5’AATGATACGGCGACCACCGAGA3’; Reverse P7-Adaptor 5’ CGTATGCCGTCTTCTGCTTG 3’) were then ligated to the synthesized fragments and subjected to end-repair processing. After agarose gel electrophoresis, 200-300 bp insert fragments were selected and used as templates for downstream PCR amplification and cDNA library preparation. The combined gill and RBC samples (1 µg RNA) were sent for RNAseq Poly-A sequencing with the Illumina NovoSeq™ 6000 platform (Novogene; Beijing, China) and raw reads are made available on NCBI (PRJNA722314).

RNAseq data was used to generate a *de novo* transcriptome assembly which was mined for white seabass isoforms of the Slc9a1 protein family using methods previously described (33). Briefly, raw reads were analyzed, trimmed of adaptor sequences, and processed with the OpenGene/fastp software (34), to remove reads: i) of low quality (PHRED quality score < 20), ii) containing >50% unqualified bases (base quality < 5), and iii) with >10 unknown bases. Any remaining unpaired reads were discarded from downstream analysis and quality control metrics were carried out before and after trimming (raw reads 80.07 x 10^6^; raw bases 12.01 Gb; clean reads 79.44 x 10^6^, clean bases 11.84 Gb, clean reads Q30 95.26%; GC content 46.67%). Thereafter, fastq files were merged into a single data set, normalized, and used for *de novo* construction of a combined gill and RBC transcriptome using the Trinity v2.6.6 software. Normalization and assembly were performed using the NCGAS (National Centre for Genome Analysis Support) *de novo* transcriptome assembly pipeline (github.com/NCGAS/de-novo-transcriptome-assembly-pipeline/tree/master/Project_Carbonate_v4) on the Carbonate High Performance Computing cluster at Indiana University. For assembly, minimum kmer coverage was set to three and the minimum number of reads needed to glue two inchworm contigs together, was set to four (35). The resulting nucleotide FASTA file was translated into six protein reading frames using BBMap (36), which were mined for the NHE-like proteins using HMMER3 v.3.0 (hmmer.org) by querying the *de novo* assembly against a hidden markov model (HHM) homology matrix generated from 132 aligned protein sequences of the vertebrate NHE family (Slc9a1 – Slc9a9; for accession numbers see Table S1 of the supplement at: doi.org/10.6084/m9.figshare.14934405.v1). Sequences were aligned using MUSCLE (37) in SeaView (38, 39), with NHE2 from *Caenorhabditis elegans* as an outgroup, and results were refined using GBlocks (40) according to the parameters specified previously (41). Phylogenetic analysis was conducted on the Cyberinfrastructure for Phylogenetic Research (CIPRES) Science Gateway (42) using the RAxML software v.8.2.12 (43) with the LG evolutionary and GTRGAMMA models (44). Branch support was estimated by bootstraps with 450 replications and the constructed tree was edited in FigTree v.1.4.4. Finally, the open reading frame of the white seabass β-NHE sequence (predicted 747 amino acids), was analyzed for the presence of the Kozak nucleic acid motifs (5’-(gcc)gccRccAUGG-3’; 45) immediately upstream of putative start codons, using the ATGpr software (46).

To confirm the expression of the β-NHE in the RBCs of white seabass, an additional blood sample of 1 ml was collected and processed as described previously. RBCs were lysed by repeatedly passing them through a 23G needle and RNA extraction was on 50 µg of RBCs by standard Trizol and chloroform extraction following the kit instructions. Isolated RNA was treated with DNAse I and 1 µg RNA was used to synthesize first-strand cDNA using SuperScript IV reverse transcriptase (Thermo Fisher 18090010). Full-length cDNA sequences were obtained in 35 cycles of PCR reactions with Phusion DNA polymerase (New England Biolabs, Ipswich, USA; MO531L) and specific primers designed against the sequence of the phylogenetically characterized white seabass β-NHE obtained from the combined gill and RBC transcriptome (Integrated DNA Technologies, Coralville, Iowa; F: 5’TCC CGT ACT ATC CTC ATC TTC A-3’ R: 5’-CCT CTG CTC TCT GAA CTG TAA AT-3’). Amplicons were analyzed by gel electrophoresis (Bio-Rad ChemiDoc) that confirmed the presence of a single band (2372 bp; Fig. S1). A-overhangs were added to Phusion products with one unit Taq polymerase (New England Biolabs; MO267S) followed by 10 min incubation at 72°C. Products were cloned (TOPO TA Cloning Kit/pCR 2.1-TOPO Vector; Invitrogen; K4500) and the ligated product was transformed into TOP10 chemically competent *E. coli* cells (Invitrogen; K457501) according to manufacturer specifications. Following over-night incubation at 37°C, single colonies of transformants were grown in Luria-Bertani (LB) broth over-night on a shaking incubator (37°C, 220 rpm; Barnstead MaxQ 4000). Plasmid DNA was isolated (PureLink Quick Plasmid Miniprep kit; Invitrogen K210010) according to manufacturer specifications and inserts were sequenced to confirm their identity and uploaded to NCBI (MW962257).

### Calculations and statistical analysis

All data were analyzed with R v.4.0.4 (47) in RStudio v.1.4.1106 (48) and figures were created with the ggplot2 package (49). Normality of the residuals was tested with the Shapiro-Wilk test (stat.desc function in R) and homogeneity of variances was confirmed with the Levene’s test (leveneTest function in R). Deviations from these parametric assumptions were corrected by transforming the raw data. All R source code is made publicly available (doi.org/10.6084/m9.figshare.14934405.v1).

To determine the blood O_2_-binding characteristics, Hb P_50_ and n_H_ values were those determined with the BOBS software v.1.0.20 (Loligo) and oxygen equilibrium curves (OEC) were generated by fitting a two parameter Hill function to the mean P_50_ and n_H_ for 8 individual fish. The main effects of PCO_2_ on P_50_ and n_H_ were analyzed with ACOVA (*P* < 0.05, *N* = 8). Plasma [HCO_3_^-^] was calculated from TCO_2_ by subtracting the molar [CO_2_] calculated from the dissociation constant and solubility coefficients in plasma at 17°C and the corresponding sample pH (6). The Bohr effects relative to pH_i_ and pH_e_, the relationship between RBC pH_i_ and pH_e_ and the non-bicarbonate buffer capacity of whole blood were determined by linear regression analysis, results of which are shown in detail in the supplement (Fig. S2A-D). The average values for these blood characteristics are shown in the main text and were calculated as the average slopes across all individuals.

In the RBC swelling trial, mean cell Hb was calculated as [Hb] divided by Hct as a decimal. Since Hb is a membrane impermeable solute, MCHC is used as a common indicator of RBC swelling. Main effects of drugs (DMSO, ISO and ISO+Am), time (10, 30 and 60 min) and their interaction (drug×time) on Hct, [Hb], MCHC, pH_e_ and pH_i_ were determined by two-way ANOVAs (lm and Anova functions in R; *N* = 5-6; *P* < 0.05) and multiple comparisons were conducted with t-tests (pairwise.t.test function in R) and controlling the false detection likelihood (FDR) with a Benjamini-Hochberg correction (p.adjust function in R).

The effect of RBC β-adrenergic stimulation on Hb-O_2_ binding was assessed by analyzing the absorbance data from the BOBS in R (R source code available at github.com/tillharter/White-Seabass-beta-NHE). In brief, the absorbances recorded at 430 nm were divided by the isosbestic wavelength of 390 nm (where absorbance is independent of Hb-O_2_ binding), and these ratios were used as the raw data for subsequent analyses. For each trace, the ten final absorbance ratios at each equilibration step were averaged (i.e. 10 s) and Hb-O_2_ saturation was calculated relative to the absorbance at the two initial calibration conditions (i.e. high: 99.7 kPa PO_2_, 0.3 kPa PCO_2_; and low: 0 kPa PO_2_, 0.3 kPa PCO_2_), assuming full saturation or desaturation of Hb, respectively. These calibration values were measured again at the end of each trial and a linear correction of drift was performed for each sample. The resulting values for Hb-O_2_ saturation were plotted against PCO_2_ and several non-linear models were fit to the data (Michaelis-Menten, Exponential and Hill). The model with the best fit (lowest AIC) was a three-parameter Hill function that was applied to each individual trace. The parameter estimates from this model yielded the maximal reduction in Hb-O_2_ saturation (Max. ΔHb-O_2_ sat.; %) in normoxia (21 kPa PO_2_) and hypoxia (3 kPa PO_2_), and the PCO_2_ at which this response was half-maximal (EC_50_PCO_2_; kPa). The main effects of drugs (DMSO, ISO and ISO+Am), O_2_ (normoxia and hypoxia) and their interaction (drug×O_2_), on the parameter estimates from the Hill functions were determined by two-way ANOVAs (lm and Anova functions in R; *N* = 6; *P* < 0.05). When significant main effects were detected, multiple comparisons were conducted with t-tests (pairwise.t.test function in R) and controlling the false detection likelihood (FDR) with a Benjamini-Hochberg correction (p.adjust function in R). The Root effect was determined from the non-linear model, as the Max. ΔHb-O_2_ sat. of the control treatment (DMSO). To quantify the relative changes in Hb-O_2_ saturation (ΔHb-O_2_ sat.) due to drug treatments, data were expressed relative to the paired measurements in the DMSO treatment for each individual fish and relative to the initial Hb-O_2_ saturation at 0.3 kPa PCO_2_ (i.e. 95.6 and 55.0% Hb-O_2_ saturation in normoxia and hypoxia, respectively). All data are presented as means±s.e.m.

## Results

### Blood O_2_-binding characteristics

The blood O_2_-binding characteristics of white seabass are summarized in Figure 1. When PCO_2_ was increased from arterial tension (0.3 kPa) to severe hypercapnia (2.5 kPa) Hb P_50_ increased significantly from 2.9±0.1 to 11.8±0.3 kPa. At the same time, the cooperativity of Hb-O_2_ binding, expressed by n_H_, decreased significantly from 1.52±0.04 to 0.84±0.03, which was reflected in a change in the shape of the OECs from sigmoidal to hyperbolic. Over the tested range of PCO_2_, white seabass displayed a Bohr coefficient of -0.92±0.13 when expressed relative to the change in pH_e_ and -1.13±0.11 when expressed relative to the change in RBC pH_i_ (Fig. S2A and B). In addition to the reduction in Hb-O_2_ affinity at elevated PCO_2_, white seabass blood also displayed a large Root effect, where the non-linear model predicted a maximal reduction of Hb-O_2_ carrying capacity of 52.4±1.8%. The relationship between pH_i_ and pH_e_ had a slope of 0.67±0.07 (Fig. S2C), reflecting the higher buffer capacity of the intracellular space. The non-bicarbonate buffer capacity of white seabass whole blood was -2.43±0.56 mmol l^-1^ pH_e_^-1^ at a Hct of 5% (Fig. S2D). By correcting this value for the higher Hct *in vivo*, according to Wood et al. (50), white seabass with a Hct of 25% are expected to have a whole blood non-bicarbonate buffer capacity of -9.68 mmol l^-1^ pH_e_^-1^.

**Figure 1.**
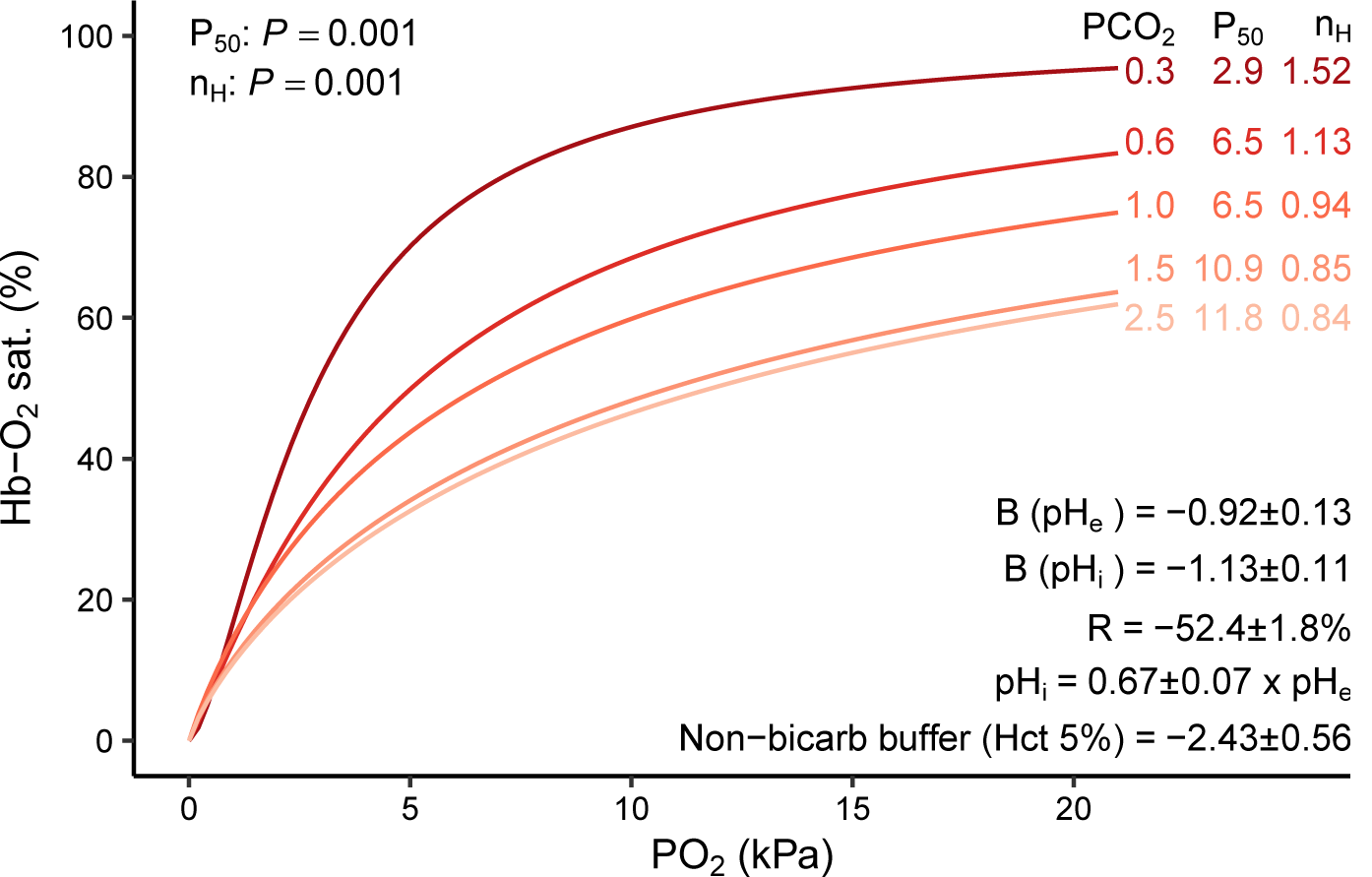
Oxygen binding characteristics of white seabass whole blood. Oxygen equilibrium curves showing hemoglobin-oxygen saturation (Hb-O_2_ sat.; %) as a function of the partial pressure of oxygen (PO_2_; kPa) at five partial pressures of carbon dioxide (PCO_2_). The PO_2_ that yields 50% Hb-O_2_ saturation (P_50_) and the cooperativity coefficient of Hb-O_2_ binding (Hill coefficient, n_H_) are shown for each curve. The main effects of PCO_2_ on P_50_ and n_H_ were analyzed with ACOVA (*P* < 0.05, *N* = 8). The Bohr coefficients (B) relative to extracellular (pH_e_) and intracellular (pH_i_), the relationship between pH_e_ and pH_i_ and the non-bicarbonate buffer capacity of the blood (at 5% Hct) were determined by linear regressions (see Fig. S2). The Root effect (R) was determined at 21 kPa PO_2_ from the model parameters shown for the DMSO treatment in Fig. 6B. All data are means±s.e.m.

### RBC swelling after β-adrenergic stimulation

The β-adrenergic stimulation of white seabass blood with ISO induced changes in the measured blood parameters (Fig. 2). Significant main effects of drug, time and their interaction (drug×time), indicate that Hct was affected by the experimental treatments (Fig. 2A). A large increase in Hct was observed in ISO-treated blood that was absent in ISO+Am and DMSO-treated RBCs. In addition, a main effect of drug treatments on MCHC indicated that the increase in Hct after ISO addition was due to swelling of the RBCs (Fig. 2B), whereas [Hb] was not affected by the treatments (Fig. S3). Significant main effects of drug and time were also detected for pH_e_, where a large extracellular acidification was observed in ISO-treated blood, relative to the DMSO and ISO+Am treatments (Fig. 2C). No significant main effect of drug or time were observed on RBC pH_i_, but multiple comparisons indicated a trend for a higher pH_i_ in ISO compared to DMSO treated blood (*P* = 0.081; Fig. 2D). Differential interference contrast (DIC) images confirmed a normal morphology of the RBCs at the beginning and the end of the trials, thus validating the fixation procedure. Swelling was observed in ISO-treated RBCs, relative to initial measurements, or DMSO and ISO+Am-treated cells (Fig. 2E-H). The swelling of ISO-treated RBCs occurred largely along the z-axis of the cells (indicated by the arrows), whereas no visible distortion was observed in the x-y directions.

**Figure 2.**
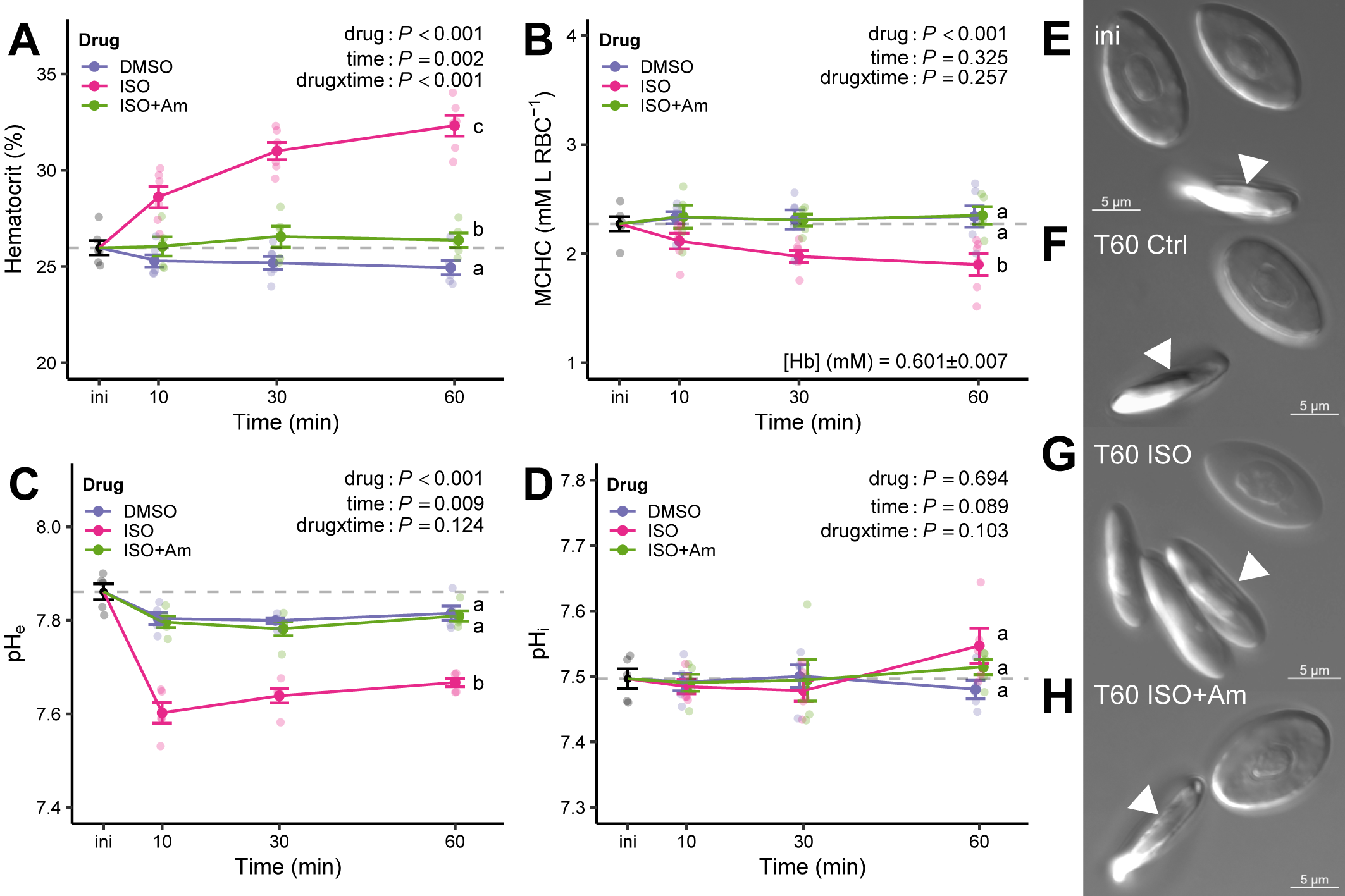
Changes in blood parameters after adrenergic stimulation of white seabass whole blood. A) Hematocrit (%), B) mean cell hemoglobin content (MCHC; mM hemoglobin l^-1^ red blood cells), C) extracellular pH (pH_e_) and D) intracellular pH (pH_i_). Blood was equilibrated in tonometers at 3 kPa PO_2_ and 1 kPa PCO_2_ and treated with either: i) a carrier control (DMSO; 0.25%), ii) the β-adrenergic agonist isoproterenol (ISO; 10 µM), or iii) ISO plus amiloride (ISO+Am; 1 mM), an inhibitor of sodium-proton exchangers (NHE). The dotted line indicates initial values for each parameter and changes were recorded over 60 min. The main effects of drug, time, and their interaction term (drug×time) were analyzed with a two-way ANOVA (*P* < 0.05, *N* = 6 and *N* = 5 for ISO+Am). There were no significant changes in hemoglobin concentration ([Hb]; mM) throughout the trials. Multiple comparisons were with paired t-tests with a Benjamini-Hochberg correction and superscript letters that differ indicate significant differences between treatments at 60 min. Individual datapoints and means±s.e.m. Inserts E-H) differential interference contrast (DIC, 60x) images of red blood cells fixed at the beginning (ini) and the end of the trial (T60). Cell swelling was visually confirmed in ISO-treated cells, but not in other treatments, and mostly along the z-axis of the cells (arrows), while the x-y-axis seemed largely unaffected.

### Molecular β-NHE characterization

The combined gill and RBC *de novo* transcriptome of white seabass contained nine Slc9a1 transcripts and phylogenetic analysis placed these sequences within well-supported clades of the NHE family tree (Fig. 3). Importantly, one of these white seabass transcripts grouped with the Slc9a1b sequences from other teleost fishes, supporting its classification as a white seabass β-NHE. Results from RT-PCR, cloning and sequencing confirmed the expression of β-NHE mRNA in isolated white seabass RBCs. A search for the Kozak nucleic acid motif in the open reading frame of the white seabass β-NHE sequence yielded the five most likely potential start codons, including one that would produce a 66 kDa protein (Table S2). This size closely matched the single band that was specifically recognized by the polyclonal β-NHE antibody in Western blots with crude homogenate, cytosolic and membrane-enriched fractions of a white seabass RBC lysate (Fig. S4A); whereas no immunoreactivity was observed in lanes where the antibody had been incubated with its pre-immune peptide. The anti-α-tubulin antibody detected a single band in the RBC crude homogenate, at the predicted size of 54 kDa (Fig. S4B). Finally, the white seabass β-NHE protein sequence shared seven consecutive amino acids with the peptide used to raise the polyclonal antibodies (Fig. S4C), which is sufficient for specific antibody binding (51). More importantly, the antigen peptide sequence was absent in the other eight white seabass NHE isoforms, ruling out non-specific antibody recognition of these NHEs. *Subcellular localization of RBC β-NHE*

**Figure 3:**
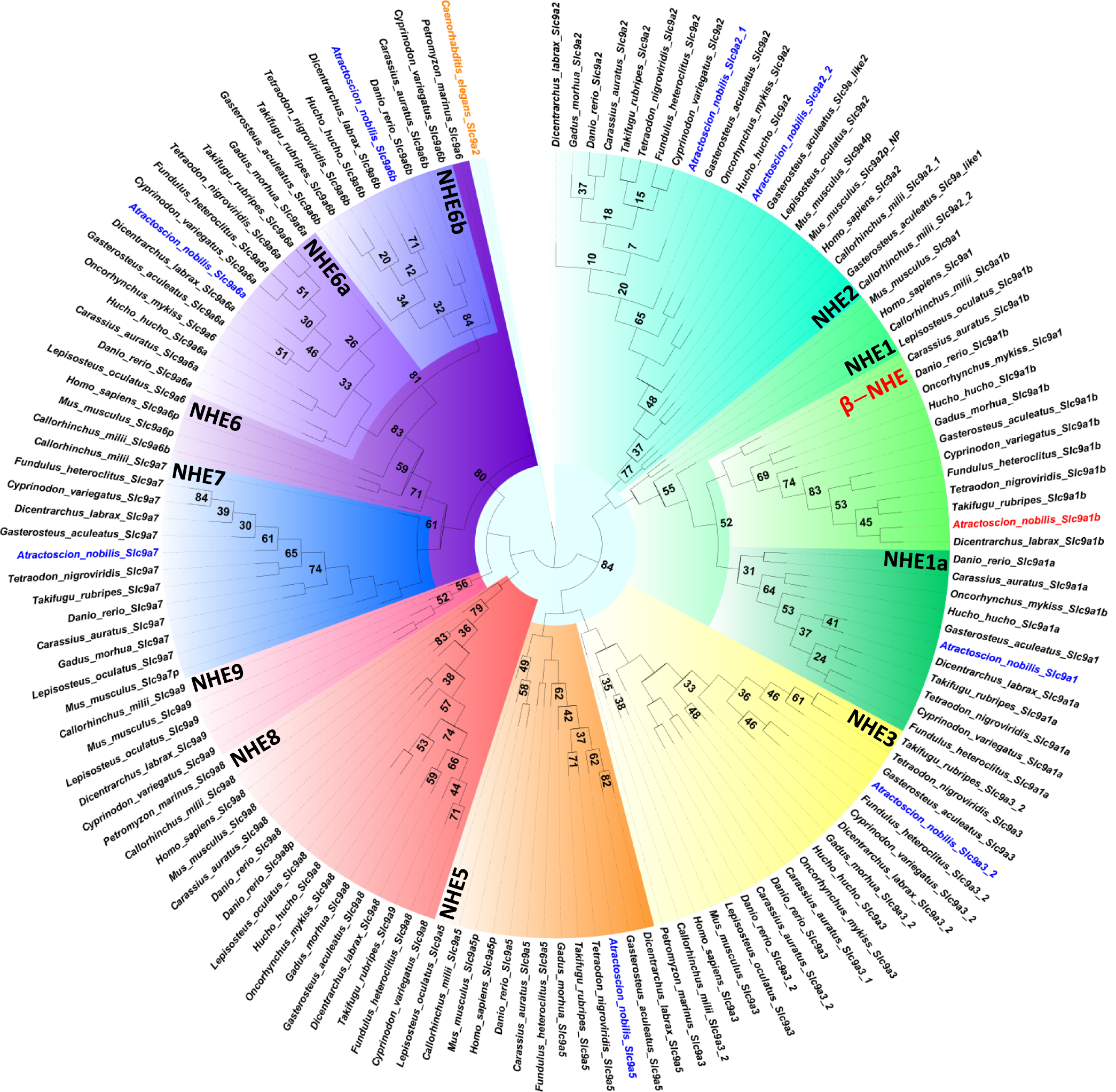
Phylogenetic analysis of nine NHE-like protein sequences in the *de novo* assembly of a combined white seabass gill and red blood cell transcriptome. Novel white seabass sequences are highlighted in blue and the β-NHE in red (Slc9a1b). Shadings delineate sub-families of the Slc9a1 gene family. The tree was rooted against the NHE2 from *Caenorhabditis elegans* in orange. Accession numbers for all species are those reported in Table S1.

The subcellular location of β-NHE protein in white seabass RBCs was determined by immunofluorescence cytochemistry and super-resolution confocal microscopy (Fig. 4). In DMSO-treated RBCs, the β-NHE immunolabelling was most intense in intracellular vesicle-like structures, and weaker at the plasma membrane. There was substantial heterogeneity in the staining pattern for β-NHE in these control cells, with varying amounts of intracellular and membrane staining. In ISO-treated RBCs, the staining pattern for β-NHE was more homogeneous and most cells showed intense immunoreactivity for β-NHE in the membrane that co-localized with α-tubulin in the marginal band, and the intracellular, vesicle-like staining that was observed in the control cells was reduced (see Fig. S5 for an overview image with more cells). In contrast, ISO-treated cells that were incubated without the primary antibody or where the antibody was pre-absorbed with its pre-immune-peptide showed no immunoreactivity for β-NHE (Fig. S6). Finally, optical sectioning and three-dimensional reconstruction of these RBCs confirmed that the membrane staining for β-NHE occurred in a single plane and co-localized with α-tubulin in the marginal band (see 3D movies S1 and S2).

**Figure 4.**
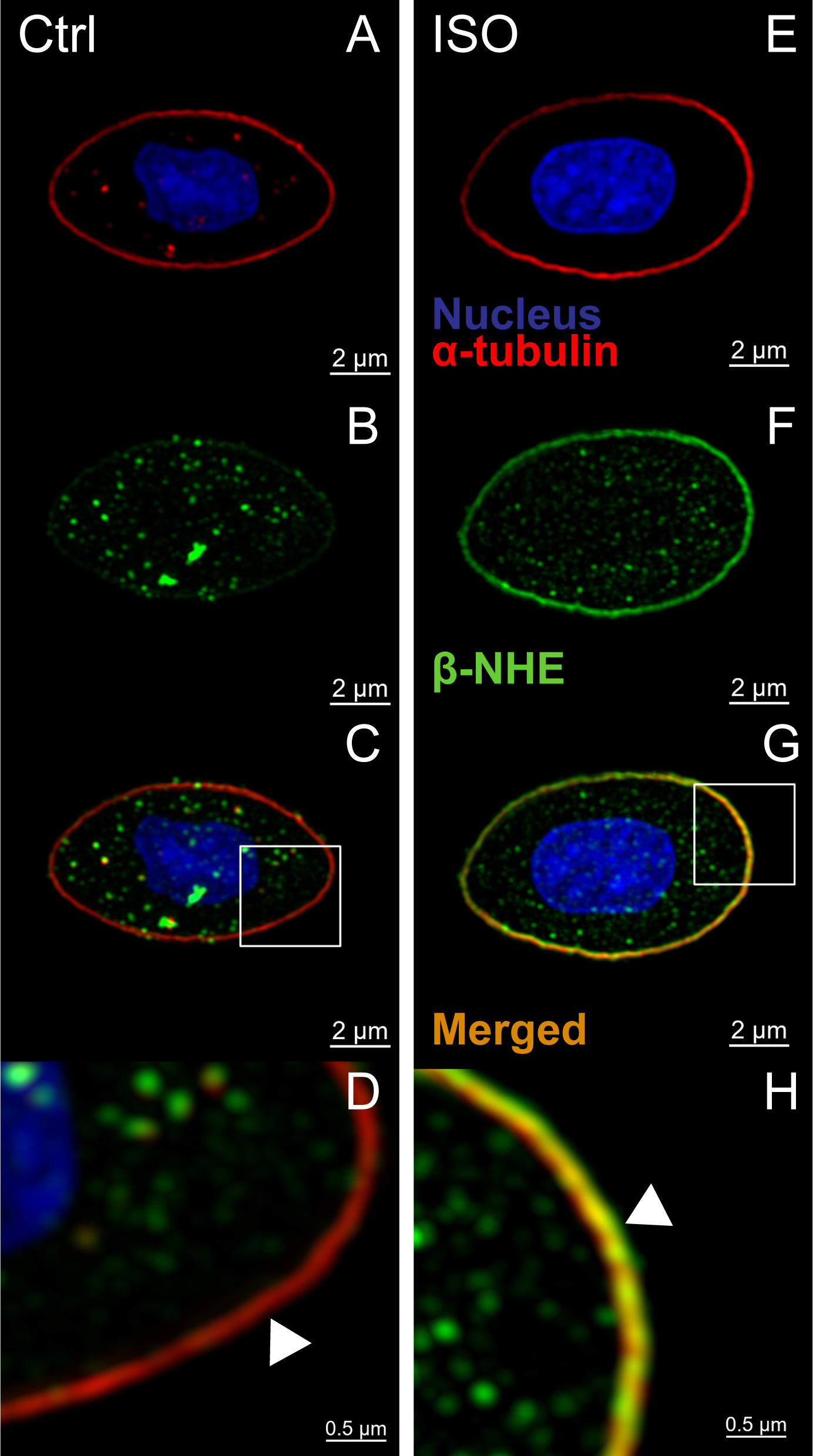
Immunocytochemical localization of the β-adrenergic sodium proton exchanger (β-NHE) in white seabass red blood cells. Blood was equilibrated in tonometers at 3 kPa PO_2_ and 1 kPa PCO_2_ for 60 mins (see Fig. 2 for details) in the presence of either: A-D) a carrier control (DMSO; 0.25%), or E-H) the β-adrenergic agonist isoproterenol (ISO; 10 µM). Fixed cells were immuno-stained with a monoclonal α-tubulin antibody to visualize the marginal band (red), with DAPI to visualize the cell nuclei (A and E), and with a polyclonal anti-β-NHE antibody (green, B and F). D and H) Magnified view of the insets in the merged images, where arrows indicate weak or absent β-NHE immunoreactivity on the membrane of Ctrl cells and intense staining in ISO-treated cells. These responses were representative and repeatable (*N* = 4) and images showing a larger number of cells are available in the supplement (Fig. S5).

### Hb-O_2_ binding after β-adrenergic stimulation

To characterize the protective effect of RBC β-NHE activation on Hb-O_2_ binding, blood samples were first equilibrated to 21 kPa PO_2_ and 0.3 kPa PCO_2_ in tonometers and no significant effects of drug treatment (DMSO, ISO or ISO+Am) were observed on any of the measured blood parameters (Fig. S7); average values were: Hct 5.20±0.14% (*P* = 0.095), [Hb] 0.178±0.006 mM (*P* =0.889), MCHC 3.46±0.15 mM l^-1^ RBC (*P* = 0.490), pH_e_ 7.848±0.018 (*P* = 0.576), pH_i_ 7.464±0.021 (*P* = 0.241). Thereafter, blood was loaded into the BOBS, where Hb-O_2_ saturation was measured spectrophotometrically at increasing levels of a respiratory acidosis in normoxia (21 kPa PO_2_). As expected from the pH-sensitivity of Hb-O_2_ binding in white seabass, an increase in PCO_2_ from 0.3 to 3 kPa caused a severe reduction in Hb-O_2_ saturation in all treatments via the Root effect (Fig. 5). The raw data were analyzed by fitting a three-parameter Hill model to the individual observations within each treatment and significant differences were observed in the parameter estimates that describe these models. EC_50_PCO_2_ was affected by the experimental treatments, as shown in a significant main effect of drug (Fig. 6A). Multiple comparisons confirmed significant differences in EC_50_PCO_2_, which was 0.85±0.06 kPa in DMSO, 0.91±0.06 kPa in ISO+Am and 1.08±0.06 kPa in ISO-stimulated blood. In contrast, the magnitude of the responses, Max. ΔHb-O_2_ sat., was not affected by the experimental treatments and no significant main effect of drug was detected; the average Max. ΔHb-O_2_ sat. across treatments was -51.1±0.7% (Fig. 6B).

**Figure 5.**
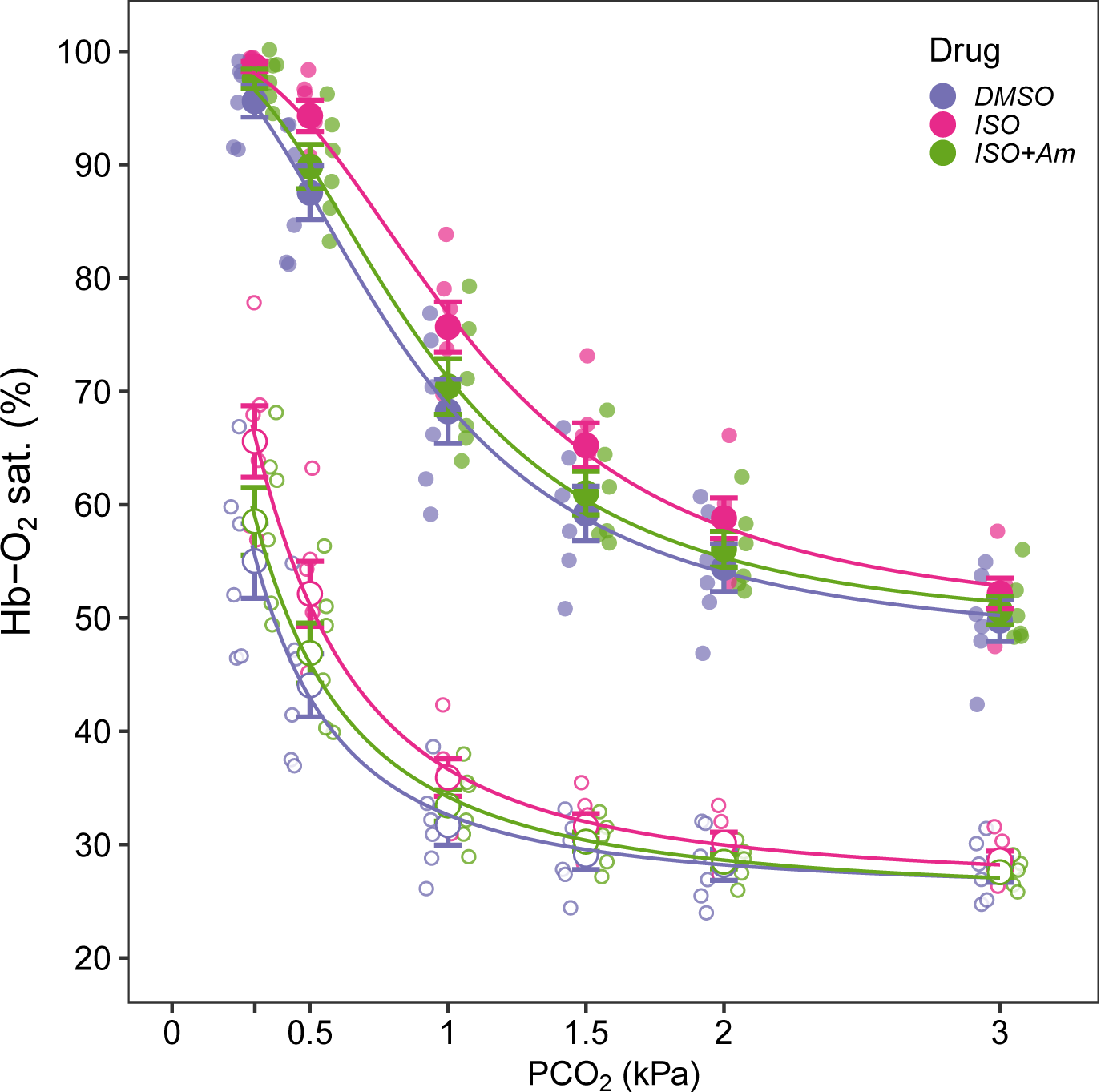
Hemoglobin-oxygen saturation (Hb-O_2_ sat.; %) during hypercapnic acidification of white seabass whole blood. Hematocrit was set to 5%, blood was equilibrated in tonometers at 21 kPa PO_2_ and 0.3 kPa PCO_2_ and treated with either: i) a carrier control (DMSO; 0.25%), ii) the β-adrenergic agonist isoproterenol (ISO; 10 µM), or iii) ISO plus amiloride (ISO+Am; 1 mM), an inhibitor of sodium-proton exchangers (NHE). For each sample, runs were performed in normoxia (21 kPa PO_2_; solid symbols) or hypoxia (3 kPa PO_2_; open symbols). Individual datapoints and means±s.e.m. (*N* = 6).

**Figure 6.**
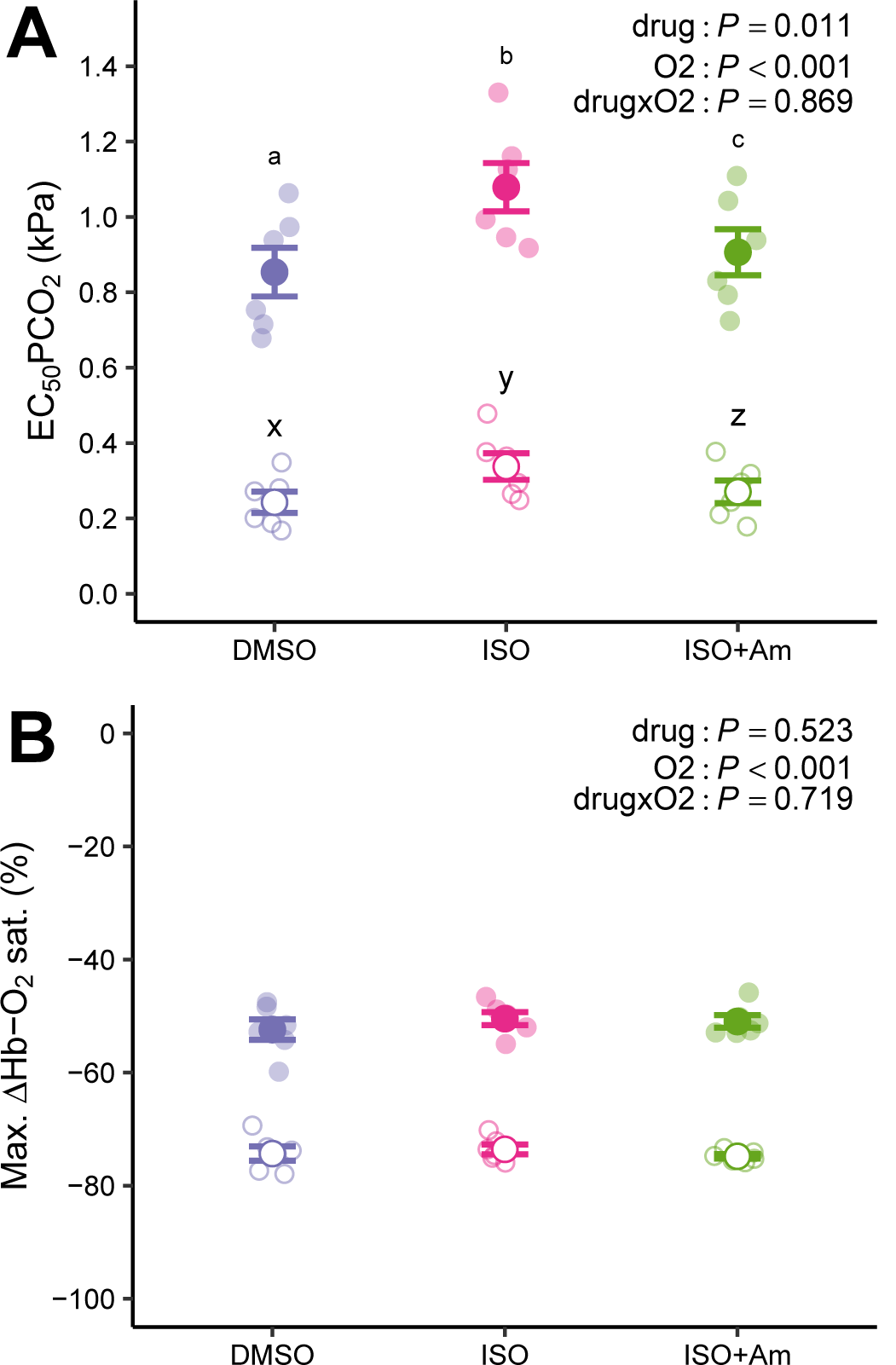
Parameter estimates describing the changes in hemoglobin-oxygen saturation during hypercapnic acidification of white seabass whole blood. A) The PCO_2_ that elicits a half-maximal reduction in Hb-O_2_ saturation (EC_50_PCO_2_; kPa). and B) the maximal reduction in Hb-O_2_ saturation due to acidification (Max. ΔHb-O_2_ sat.; %). Treatments were: i) a carrier control (DMSO; 0.25%), ii) the β-adrenergic agonist isoproterenol (ISO; 10 µM), or iii) ISO plus amiloride (ISO+Am; 1 mM) an inhibitor of sodium-proton exchangers (NHE). For each sample, runs were performed in normoxia (21 kPa PO_2_; solid symbols) or hypoxia (3 kPa PO_2_; open symbols). The main effects of drug treatments (drug), oxygen (O_2_) and their interaction term (drug×O_2_) were analyzed with a two-way ANOVA (*P* < 0.05, *N* = 6). Multiple comparisons were with paired t-tests and a Benjamini-Hochberg correction and superscript letters that differ indicate significant differences between treatments for each O_2_ tension. Individual datapoints and means±s.e.m. (*N* = 6).

When the same experiment was repeated under hypoxic conditions (3 kPa PO_2_), an increase in PCO_2_ likewise caused a severe reduction in Hb-O_2_ saturation, indicating that in white seabass, a Root effect can also be expressed at saturations around P_50_ (Fig. 5). A significant main effect of O_2_ indicated that cells in the hypoxic condition required a lower EC_50_PCO_2_ to achieve Max. ΔHb-O_2_ sat., compared to the normoxic condition (Fig. 6A). There was also a significant main effect of drug on EC_50_PCO_2_ and multiple comparisons indicated a similar pattern in the individual drug effects as in the normoxic experiment, which was further confirmed by the absence of a significant drug×O_2_ interaction. Finally, a significant effect of O_2_ on Max. ΔHb-O_2_ sat. (Fig. 6B) indicated a larger response magnitude in hypoxic blood, but that was unaffected by drug treatments, and the average Max. ΔHb-O_2_ sat. across treatments was -74.1±0.1%.

To quantify the protective effect of RBC β-NHE activation on Hb-O_2_ binding during a hypercapnic acidosis, Hb-O_2_ saturation was expressed relative to the paired measurements in the DMSO treatment for each individual fish and relative to the initial Hb-O_2_ saturation at 0.3 kPa PCO_2_ (i.e. 95.6 and 55.0% Hb-O_2_ saturation in normoxia and hypoxia, respectively). In normoxia, the benefit of β-NHE stimulation with ISO showed a bell-shaped relationship with a maximal ΔHb-O_2_ sat. of 7.8±0.02% at 1 kPa PCO_2_ (Fig. 7A). When NHEs were inhibited in ISO+Am blood, ΔHb-O_2_ sat. was only 1.9±0.4% at 1 kPa PCO_2_ and significantly lower compared to the other treatments; at higher PCO_2_ the 95% confidence intervals overlapped with the DMSO values, indicating no difference from controls. In hypoxic blood, the ISO treatment had the largest effects on ΔHb-O_2_ sat. at 0.3 kPa PCO_2_, with maximal values of 19.2±0.0% that decreased towards higher PCO_2_ (Fig. 7B). Whereas, in the ISO+Am treatment, ΔHb-O_2_ sat. was 6.4±0.0% at 0.3 kPa PCO_2_, and significant differences to the DMSO controls were only observed at PCO_2_ below 1.5 kPa.

**Figure 7.**
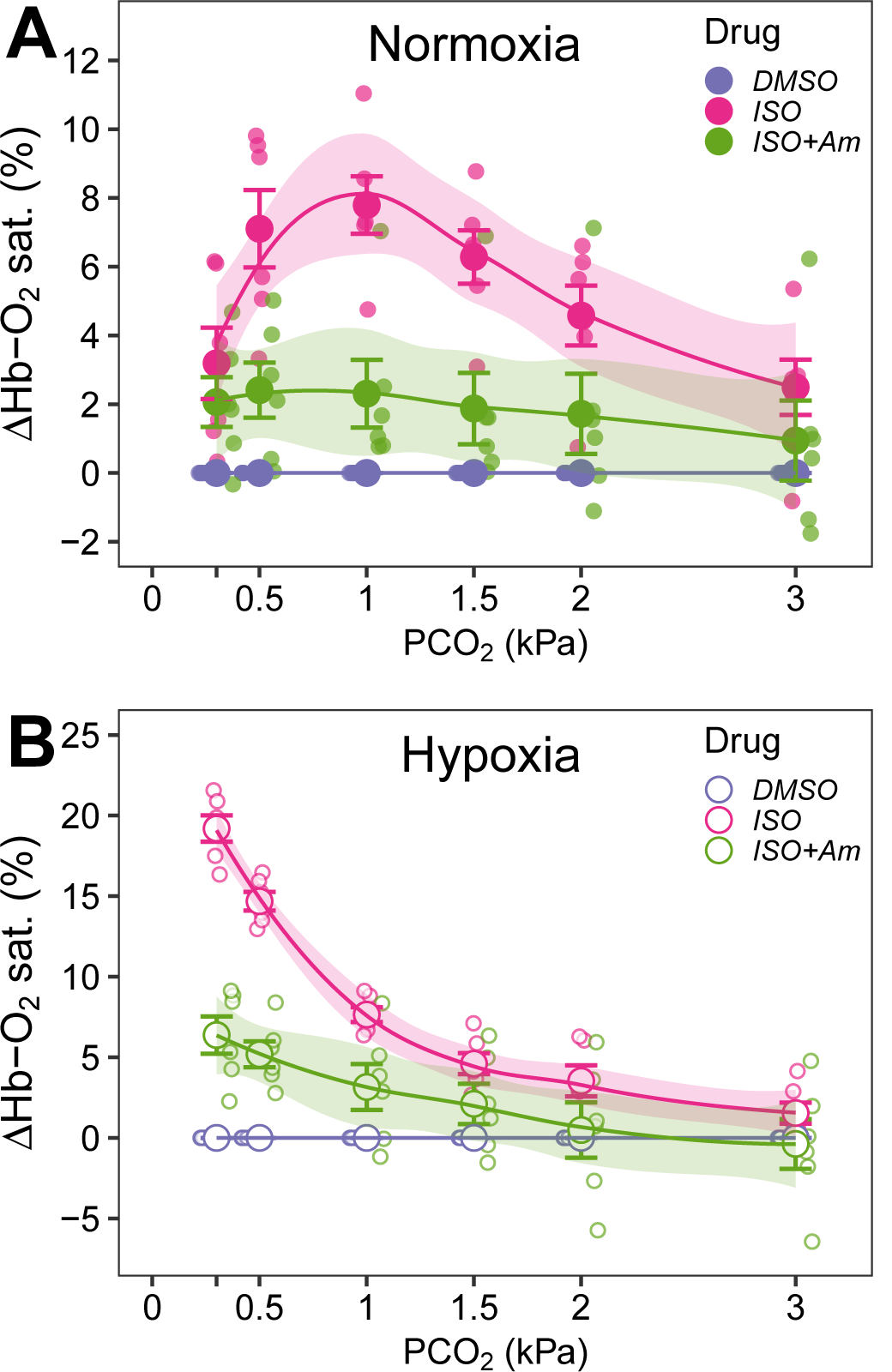
Relative changes in hemoglobin-oxygen saturation (ΔHb-O_2_ sat.; %) during hypercapnic acidification of white seabass whole blood. A) in normoxia (21 kPa PO_2_; solid symbols) or B) hypoxia (3 kPa PO_2_; open symbols). Treatments were: i) a carrier control (DMSO; 0.25%), ii) the β-adrenergic agonist isoproterenol (ISO; 10 µM), or iii) ISO plus amiloride (ISO+Am; 1 mM), an inhibitor of sodium-proton exchangers (NHE). Individual datapoints, means±s.e.m. and 95% confidence intervals (*N* = 6).

## Discussion

In line with our initial hypothesis, RBC β-NHE activity in white seabass may greatly protect the blood O_2_-carrying capacity during environmentally relevant levels of hypoxia and hypercapnia. However, not all predictions were met as expected: white seabass did not have an unusually high Hb-O_2_ affinity and thus, other aspects of their physiology are likely more important in determining their tolerance to hypoxia. Like other teleosts, white seabass had highly pH-sensitive Hbs, where a reduction in pH decreased both Hb-O_2_ affinity via the Bohr effect and Hb-O_2_ carrying capacity via the Root effect. Several lines of evidence corroborated the presence of a RBC β-NHE in white seabass and super-resolution imaging revealed, for the first time, the subcellular location of β-NHE protein in intracellular, vesicle-like structures and on the RBC membrane. Furthermore, adrenergic stimulation induced changes in the intracellular distribution of the β-NHE that may indicate a role of protein translocation in regulating β-NHE activity. A detailed quantification of the protective effects of RBC β-NHE activity, revealed the largest benefits at ∼1 kPa PCO_2_ in normoxia (21 kPa PO_2_), where Hb-O_2_ saturation increased by ∼8%. Whereas in hypoxia (3 kPa PO_2_), β-NHE activity had its largest effect at arterial PCO_2_ (0.3 kPa) and enhanced Hb-O_2_ saturation by ∼20%; however, the benefits of β-NHE activation in hypoxia decrease rapidly at higher PCO_2_, revealing a potential vulnerability of white seabass to combinations of these stressors.

Many hypoxia tolerant vertebrates have evolved Hbs with a high affinity for O_2_ (low Hb P_50_ values), which helps to extract the gas from the respiratory medium (9, 52, 53). White seabass in the present study had a Hb P_50_ of 2.9±0.1 kPa (Fig. 1), which is higher than the values typically found in hypoxia tolerant fishes, such as carp (*Cyprinus carpio*) that have Hb P_50_ values as low as 0.5 kPa (54). In fact, the Hb P_50_ of white seabass resemble more closely the values in the well-studied rainbow trout, of 3.3 kPa (55), a cold-stream salmonid, of no noteworthy hypoxia tolerance. However, the O_2_-binding affinity of Hb must strike a balance between loading O_2_ at the gas exchange surface and unloading O_2_ at the tissues (56). Everything else being equal, a higher Hb P_50_ can sustain a higher PO_2_ at the tissue capillaries, enhancing the diffusion gradient of O_2_ to the mitochondria, which is of particular benefit to those species with a high scope for exercise (57). Thus, it seems that a high Hb-O_2_ affinity is not part of the physiological mechanism that facilitates hypoxia tolerance in white seabass, but instead, a high tissue PO_2_ may be important to sustain exercise performance in these active piscivores.

As in other teleosts, especially those in the highly-derived group of perciformes, Hb-O_2_ binding in white seabass was highly pH-sensitive. An increase in PCO_2_ from arterial levels (0.3 kPa) to severe hypercapnia (2.5 kPa), caused a significant right-shift of the OEC (Fig. 1) via the Bohr effect, increasing P_50_ to 11.8±0.3 kPa. When considering the corresponding changes in pH_e_ (from 8.1 to 7.2 over the range of tested PCO_2_; see Fig. S2A) the Bohr coefficient in white seabass was -0.92, and slightly higher, at -1.13, when considering the changes in RBC pH_i_ (from 7.7 to 7.0: Fig. S2B). Again, these Hb-O_2_ binding characteristics resemble closely those of rainbow trout, where P_50_ increases to 10 kPa at 2 kPa PCO_2_, yielding a Bohr coefficient (relative to pH_e_) of -0.87 (55). A normoxic increase in PCO_2_ caused a significant reduction in Hb-O_2_ saturation via the Root effect, and at PCO_2_ above 3 kPa the O_2_-carrying capacity of white seabass Hb was reduced by 52.4±1.8% (DMSO treatment; Fig. 6B). These results are in line with those of other teleosts, such as rainbow trout (∼55%), tench (*Tinca tinca*; ∼50%) and the European perch (*Perca fluviatilis*; ∼70%), where the larger Root effect values may reflect the higher final PCO_2_ used during those trials (58).

The Root effect is part of a specialized system of O_2_ supply to the eye and the swimbladder of teleosts, where blood is acidified in a counter-current exchanger (the *rete mirabile*) to produce high PO_2_ that bridge the large diffusion distances to the avascular retina of teleosts and inflate the swimbladder against large hydrostatic pressures (59). In the course of teleost evolution there have been numerous secondary losses of the *choroid* and swimbladder *retes*. While their presence has not directly been determined in white seabass, an ancestral state reconstruction predicts no secondary loss of either *rete* on the teleost branch leading up to the perciformes, which include the white seabass (24). In addition, all of the five independent losses of the *choroid rete* have coincided with a reduction of the Root effect below 40% (24, 60). Thus, the large Root effect of white seabass is consistent with the presence of a *choroid rete* and likely critical for maintaining a high ocular PO_2_ that facilitates the visual acuity in these active predators (15).

Vertebrate Hbs are intracellular proteins and, as such, are affected by the microenvironment within the RBC cytoplasm. Teleost β-NHEs can actively modulate Hb-O_2_ binding by controlling RBC pH_i_ and several lines of evidence in our study indicate the presence of functional β-NHEs in white seabass RBCs. A combined gill and RBC transcriptome detected nine sequences belonging to the vertebrate NHE (Slc9a1) family and phylogenetic analysis classified the white seabass Slc9a1b transcript as belonging to the larger group of teleost RBC β-NHEs (Fig. 3). These findings were supported by the results from RT-PCR, confirming the expression of the β-NHE in white seabass RBCs. Western blots with a polyclonal anti-trout β-NHE antibody recognized a single band at 66 kDa in white seabass RBC homogenates (Fig. S4A), which is smaller than the 84 kDa predicted based on the longest possible mRNA transcript (Table S2). However, a search for Kozak motifs revealed the five most likely potential start codons in the open reading frame of the white seabass β-NHE mRNA sequence, one of which predicted a protein size of 66 kDa that matches the protein size detected in Western blots. This predicted β-NHE isoform lacks 158 amino acids on the N-terminus, which, according to structural NHE-protein models (61), are not essential for the transporter’s activity, but may determine a differential sensitivity to inhibitors (61, 62). NHE isoforms from other teleosts have also been shown to separate in Western blotting with a similar size discrepancy (63), and show differential sensitivity to amiloride and its derivatives compared to mammalian NHEs (64).

Adrenergic stimulation of white seabass RBCs with ISO caused a ∼25% volume increase during the 60 min trials, whereas no changes in RBC volume were detected in DMSO treated cells. The swelling response was corroborated by a significant reduction in MCHC and by visually confirming the increase in cell volume under a microscope, and these results closely match previous reports of RBC swelling after adrenergic stimulation in other teleosts (23, 65, 66). In addition, the ISO-induced swelling was abolished by the inhibition of NHEs in ISO+Am-treated RBCs, providing additional pharmacological support for the presence of a RBC β-NHE in white seabass. In ISO-treated RBCs, but not those treated with DMSO or ISO+Am, we observed a decrease in pH_e_, which is the direct result of H^+^ excretion by NHE activity. Corresponding changes in RBC pH_i_ are typically smaller, due to the higher buffer capacity of the intracellular space (pH_i_ = 0.67±0.07×pH_e_; Fig. 1), and the additional freeze-thaw steps and plasma removal increase the variability of these measurements. Consequently, we were not able to resolve significant treatment effects on RBC pH_i_, but a non-significant trend may point towards a small increase in RBC pH_i_. Another interesting observation in these RBC swelling trials were the changes in cell morphology due to adrenergic stimulation. The increase in cell volume was largely due to an expansion along the z-axis of the cells, whereas the dimensions in the x-y axis apparently remained unaffected. The nucleated RBCs of non-mammalian vertebrates, including fish, have a marginal band, a structural component of their cytoskeleton formed by strands of α-tubulin that maintains their elliptical shape in the face of shear and osmotic disturbances (67). The stiffness of this marginal band (68) may be a major impediment to swelling along the x-y axis forcing the cells to widen in the z-direction.

Fixed RBCs from these swelling trials were studied in more detail by super-resolution microscopy and by immunolabelling β-NHE and α-tubulin (Fig. 4). All RBCs showed β-NHE immunoreactivity, corroborating the presence of β-NHE protein in these cells. In control RBCs, β-NHE protein was detected intracellularly and appeared to be confined to vesicles, while weaker staining was detected on the plasma membrane (Fig. 4B and S5). A similar staining pattern has been described for a NHE1-like protein in the RBCs of winter flounder (*Pseudopleuronectes americanus*; Pedersen et al., 2003). However, the immunolabelling of this NHE was with polyclonal antibodies raised against a region of the human NHE1 sequence (aa 631-746) that is highly conserved with both the teleost Slc9a1a and Slc9a1b. Therefore, these previous results likely include staining of several NHE isoforms including the flounder β-NHE. The antibody used in the present study showed a high specificity for the white seabass β-NHE (Fig. S4) and a confounding detection of other RBC NHE isoforms is unlikely.

An important finding of our work was that the intracellular localization of β-NHE protein changed after adrenergic stimulation of the RBCs. In ISO-treated cells the staining pattern for β-NHE was more homogeneous compared to controls, with strong signal at the plasma membrane, and weaker intracellular signal (Fig. 4F and S5). Optical sectioning and 3D reconstructions of these cells clearly showed that the intense membrane staining for β-NHE was confined to a single plane, colocalizing with α-tubulin in the marginal band, and that this staining was mostly absent in DMSO-treated cells (Movies S1 and S2). Furthermore, the use of super-resolution microscopy allowed us to discern the subcellular orientation the β-NHE signal, which was extracellular relative to α-tubulin (Fig. 4H), thus, indicating a direct contact with the blood plasma that is essential for regulating pH_i_ via NHE activity. Combined, these observations may point towards an adrenergically-induced translocation of β-NHE protein from the cytoplasm into the membrane of white seabass RBCs. Intracellular translocation of NHEs in response to various stimuli has been reported in other systems, such as the gills of acid infused hagfish (70), insulin-treated rat cardiomyocytes (71), isolated mammalian cells after acidification (72) or the initiation of Na^+^-glucose co-transport in intestinal epithelial cells (73). Studies on rainbow trout RBCs found that the abundance of radio-labelled β-NHE protein in the membrane increased after hypoxic incubation (1.2 kPa for 30 mins, 74), which generally supports a mechanism of protein translocation. However, we incubated all RBCs in hypoxia and hypercapnia, and observed β-NHE translocation after adrenergic stimulation. Clearly, there are still many open questions regarding the well-studied β-NHE response of teleosts, and additional work is required to characterize the cellular mechanisms underlying the translocation of β-NHE protein and its regulation by catecholamines, PO_2_, PCO_2_ or pH. If substantiated, these findings may open new avenues in the research of RBC pH_i_ regulation in teleosts and perhaps other vertebrates.

The activation of RBC β-NHEs has been shown to raise pH_i_ above the equilibrium condition and plays an important role in protecting Hb-O_2_ binding in teleosts (21). Previous work has characterized the resulting left-shift in the OEC (75–77) and the changes in arterial O_2_-carrying capacity due to adrenergic stimulation of the RBCs (78–80). In the present study we quantified the protective effect of β-NHE activation on Hb-O_2_ saturation in white seabass under environmentally relevant levels of hypercapnia and hypoxia. As expected, Hb-O_2_ saturation decreased significantly, due to the Root effect, when PCO_2_ was increased from 0.3-3 kPa (Fig. 5). Adrenergic stimulation of the RBCs with ISO significantly delayed the reduction in Hb-O_2_ saturation to higher EC_50_PCO_2_ that were 1.08±0.06 kPa in ISO compared to 0.85±0.06 kPa in DMSO-treated blood (Fig. 6A). In ISO+Am-treated RBC, the EC_50_PCO_2_ decreased significantly to 0.91±0.06%, compared to ISO-treated cells, corroborating the involvement of the RBC β-NHE in the response. However, the EC_50_PCO_2_ of ISO+Am-treated RBCs was still significantly higher compared to DMSO controls, perhaps indicating that 1 mM amiloride did not lead to a full inhibition of the β-NHE under the tested conditions, or that other, amiloride insensitive transporters, play a role in elevating RBC pH_i_ after adrenergic stimulation.

While β-NHE activity shifted the reduction in Hb-O_2_ saturation to a higher PCO_2_, the magnitude of the Root effect was not affected by adrenergic stimulation (Fig. 6B). No significant differences were observed in Max. ΔHb-O_2_ sat. in any of the tested treatments and therefore, a severe acidosis generated by high PCO_2_ can overwhelm the physiological capacity of the β-NHE to protect RBC pH_i_. The H^+^ extrusion by the β-NHE is secondarily active and driven by the trans-membrane Na^+^ gradient created by the RBC Na^+^-K^+^-ATPase (NKA). While both NKA activity (81) and the RBC rate of O_2_ consumption (ṀO_2_) increase after adrenergic stimulation (82), it is possible that the capacity of the NKA to maintain the larger Na^+^ gradients required to compensate for a greater reduction in pH_i_ is limited, as could be the availability of ATP to fuel the exchange. In addition, H^+^ that are extruded by the β-NHE will react with HCO_3_^-^ in the plasma to form CO_2_ that can, once again, diffuse into the cells. This re-acidification of the cells via CO_2_ is part of the Jacobs-Stewart cycle and typically rate-limited by the formation of CO_2_ in the plasma of teleosts (83). However, as pH_e_ decreases, the pool of plasma H_2_CO_3_ becomes larger (84), accelerating the Jacobs-Stewart cycle and the re-acidification of the cells, which may explain, in part, why β-NHE activity is ineffective at very high PCO_2_.

The benefit of β-NHE activity on Hb-O_2_ saturation was non-linear over the range of PCO_2_ tested, and in normoxia the bell-shaped response had a maximum at ∼1 kPa PCO_2_ (Fig. 7A). The observed relationship is likely dependent on the sigmoidal shape of the OEC, where a left-shift due to β-NHE activity has only marginal effects when Hb-O_2_ saturation is high and the curve is flat (85). In addition, β-NHE activity in many teleosts is stimulated by high intracellular [H^+^] and inhibited by high extracellular [H^+^] as pH_e_ decreases, yielding a bell-shaped relationship between β-NHE activity and pH (86). The ecological implications are noteworthy, as the protective effect of β-NHE activity on Hb-O_2_ binding is greatest over the range of PCO_2_ that wild white seabass are currently experiencing during severe red-tide or upwelling events. The increase in Hb-O_2_ saturation at these PCO_2_ is ∼8%, and the effect can be harnessed continuously with every pass of the RBCs through the gills. Everything else being equal, an increase in arterial O_2_ content can sustain a proportionally higher ṀO_2_, increasing the scope for activity or reducing the requirements for anaerobic pathways of ATP production that can lead to a toxic accumulation of metabolic by-products, such as lactate and H^+^. Thus, for fish that experience a potentially life-threatening surge in PCO_2_, an 8% increase in arterial O_2_ content could make the difference between escaping into less-noxious waters or perishing in the attempt.

In the hypoxic trials, DMSO treated blood at arterial PCO_2_ (0.3 kPa) had a Hb-O_2_ saturation of 55.0±3.3%, which was close to the target value around Hb P_50_ (Fig. 5). As in normoxia, an increase in PCO_2_ caused a significant reduction in Hb-O_2_ saturation, indicating the presence of a Root effect in hypoxia, which further decreased Hb-O_2_ saturation, even below the level of the maximal normoxic Root effect. Consequently, H^+^ binding to Hb must occur over nearly the entire range of the OEC, which stands in contrast to previous findings in rainbow trout where the Bohr effect and H^+^ binding to Hb occurred largely in the upper half of the OEC (87, 88). The possibility of inter-specific differences in the interaction between Hb-O_2_ and H^+^ binding cannot be resolved from the present data. However, it seems more likely that the kinetics of H^+^ binding that induce the Bohr effect are different from those of the Root effect, which is supported by previous work indicating different molecular mechanisms for the two effects (89, 90). The interacting kinetics of O_2_ and H^+^ binding to the Root effect Hbs of teleosts remain a worthwhile avenue for future research and studying a broader range of environmental and metabolic scenarios, in more species, may strengthen the important ecological implications of the present work.

As in normoxic blood, adrenergic activation of the β-NHE in hypoxia, increased Hb-O_2_ saturation during a hypercapnic acidosis. This protective effect of the β-NHE was reflected in a significantly higher EC_50_PCO_2_ in ISO- compared to DMSO or ISO+Am-treated RBCs (Fig. 6A). A significant main effect of O_2_ on EC_50_PCO_2_, would indicate that in hypoxic blood a lower PCO_2_ is required to desaturate Hb, compared to normoxic blood. However, this parameter estimate is influenced by the combined effects of PO_2_ and PCO_2_ on Hb-O_2_ saturation (by taking into account the full scale from 0-100%), which is not easily untangled statistically. Importantly, there was no drug×O_2_ interaction, indicating that the effect of the drugs was similar under normoxia and hypoxia, highlighting the benefit of β-NHE activation under both conditions.

In hypoxia, the benefit of β-NHE activation on Hb-O_2_ saturation was also non-linear over the tested range of PCO_2_ (Fig. 7B). β-NHE activation in hypoxic blood caused the largest increase in Hb-O_2_ saturation at arterial PCO_2_ (0.3 kPa) and the benefits decreased markedly towards higher levels of hypercapnia; likely due to the flattening of the OEC at low Hb-O_2_ saturations and perhaps some inhibition of the transporter by the increasing extracellular [H^+^]. The effect of β-NHE activity on Hb-O_2_ binding was larger in hypoxia compared to normoxia, and at 0.3 kPa PCO_2_ Hb-O_2_ saturation increased by 11±0.4%. This effect is even greater when considering that the available O_2_-carrying capacity is lower in hypoxia and when expressed relative to the available Hb-O_2_ saturation (55% in DMSO treated blood), the relative benefit of β-NHE activity was 19.2±0.0%. Many teleost β-NHEs are O_2_-sensitive (91) and the larger effects of β-NHE activity on Hb-O_2_ saturation in hypoxia may be related to a partial inhibition of the transporter in normoxia; whereas, the effect appears to be less severe in white seabass compared to other species (30, 66, 92). The nearly 20% increase in Hb-O_2_ saturation due to β-NHE activity is of great ecological significance and could be a principal pathway to safeguard arterial O_2_ transport and facilitate hypoxic survival of white seabass in the wild. However, the present data also indicate a diminishing benefit of the β-NHE response when PCO_2_ increases; thus, revealing a potential vulnerability of white seabass to the combined stressors of hypoxia and hypercapnia; surviving these conditions likely requires additional behavioral and metabolic adjustments, that are yet to be determined.

### Conclusion

The present results provide a thorough characterization of the Hb-O_2_ binding system of white seabass, a non-model marine teleost with great ecological and economic importance in Southern California. Several lines of evidence confirmed the presence of a RBC β-NHE and super-resolution microscopy may point towards a regulation of the transporter’s activity via intracellular translocation, a potentially novel pathway that deserves a more thorough investigation. In white seabass, the activity of the RBC β-NHE may provide significant protection of Hb-O_2_ binding during hypercapnic conditions with maximal benefits around the ecologically relevant level of ∼1 kPa PCO_2_. Large benefits of β-NHE activation were also observed in hypoxia, however, with a greater sensitivity to increases in PCO_2_. Combined, these data indicate that RBC function plays a critical role in modulating the O_2_-binding characteristics of the pH-sensitive Hbs in white seabass and is likely part of the suite of physiological responses that determines their hypoxia and hypercapnia tolerance. Finally, these results also highlight a potential vulnerability of white seabass to combinations of these stressors and further research is needed to study the implications for wild fish conservation along the steadily warming and eutrophicated California coast and in high density aquaculture.

## Supporting information

Movie S1 Ctrl

Movie S2 ISO

## Acknowledgements

Thanks are due to Mark Drawbridge and the Hubbs SeaWorld Research Institute (HSWRI) for generously providing the white seabass and Phil Zerofski, Jessica Hallisey, Garfield Kwan and Daniel Jio for their help with animal care.

## Endnotes

The supplemental materials are available through figshare (doi.org/10.6084/m9.figshare.14934405.v1) as well as all raw data and R source code (doi.org/10.6084/m9.figshare.14944293.v1), and sequence data is available through NCBI (see detailed accession numbers in manuscript and supplement Table S1).

