## Supplemental Materials for "Quantifying the potential for red blood cell β-adrenergic sodium-proton exchangers to protect oxygen transport in hypoxic and hypercapnic white seabass"

**For Research Article:**

**Contents:**

Supplementary Tables

Supplementary Figures

Supplementary Movies

### Supplementary Tables

**Table S1 Accession information for the Slc9a1 (NHE) isoforms that were used as a reference to mine the combined gill and red blood cell transcriptome of white seabass.**

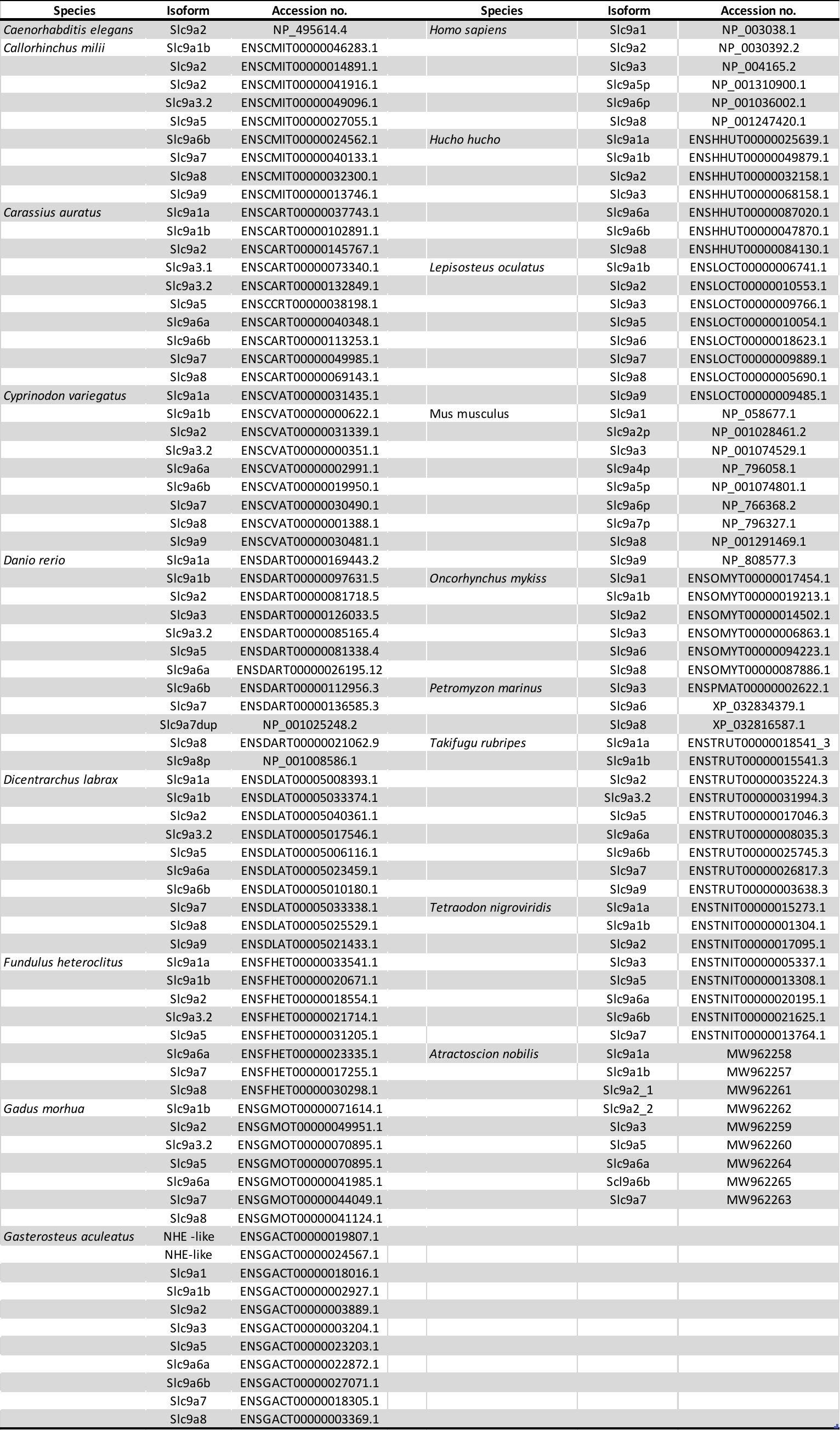

**Table S2 Kozak analysis for the five most likely start codons in the open reading frame of the white seabass β-NHE sequence detected in the combined gill and RBC transcriptome**

| **No. of ATG from 5'end** | **Reliability** | **Frame** | **Identity to Kozak rule A/GXXATGG** | **Start (bp)** | **Finish (bp)** | **ORF Length (aa)** | **Stop codon found?** | **Sequence** | **Predicted Molecular weight (kDa)** |
| --- | --- | --- | --- | --- | --- | --- | --- | --- | --- |
| 1 | 0.69 | 1 | tXXATGG | 1 | 2241 | 747 | Yes | MAVLLRPPRRSSSPLTALALRSMCVLLLVFVCAASAASRDAVEENNDTSH  HQTNSTAHKKAFPVLSFNYDHVRKPFEISLWILLALLMKLGFHIIPTVST  VVPESCLLIFVGLLVGGIIKAIGEEAPILDSKLFFLYLLPPIILDAGYFL  PIRAFTENMGTILVFAVVGTLWNAFFIGGMMYGVCQIEGAKLANVDLLSC  LLFGSIISAVDPVAVLAVFEEIHINELLHILVFGESLLNDAVTVVLYHLF  KEFSQAGTVTVVDAVLGVVCFFVVSLGGVMVGAIYGLLGAFTSRFTSHTR  VIEPLFVFLYSYMAYLSAEVFHLSGIMSLIACGVMMRPYVEANISHKSYT  TIKYFLKMWSSVSETLIFIFLGVSTVAGPHAWNWTFVVSTVVLCLVSRVL  GVIGLTFIINKFRIVKLTKKDQFIVAYGGLRGAIAFSLGFLLTNNEMKHL  FLTAIITVIFFTVFVQGMTIRPLVELLAVKRKKENKGSINEEIHTQFLDH  LLTGIECICGHYGHHHWKDKLNRFNKAYVKKWLIAGERSTEPQLISFYNK  MEMKQAMMMVESGSAAKLPTIVSSVSMQNIQPRGPARRRAIPSISKSREE  EIRKILRNNLQKTRQRLRSYSRHDLMIDPFEDNVSEIRFRKQRVEMERRM  SHYLTVPANRQETPPVRKVCFEPEHQVYTYDDSESTRGPTAQPNLSPADA  VSLLNESPQRPSQRGDGELRAKDEQEELKLSRCLSDPGPNKEEEGDG | 83.7 |
| 13 | 0.53 | 1 | AXXATGt | 979 | 2241 | 421 | Yes | MSLIACGVMMRPYVEANISHKSYTTIKYFLKMWSSVSETLIFIFLGVSTV  AGPHAWNWTFVVSTVVLCLVSRVLGVIGLTFIINKFRIVKLTKKDQFIVA  YGGLRGAIAFSLGFLLTNNEMKHLFLTAIITVIFFTVFVQGMTIRPLVEL  LAVKRKKENKGSINEEIHTQFLDHLLTGIECICGHYGHHHWKDKLNRFNK  AYVKKWLIAGERSTEPQLISFYNKMEMKQAMMMVESGSAAKLPTIVSSVS  MQNIQPRGPARRRAIPSISKSREEEIRKILRNNLQKTRQRLRSYSRHDLM  IDPFEDNVSEIRFRKQRVEMERRMSHYLTVPANRQETPPVRKVCFEPEHQ  VYTYDDSESTRGPTAQPNLSPADAVSLLNESPQRPSQRGDGELRAKDEQE  ELKLSRCLSDPGPNKEEEGDG | 48.2 |
| 22 | 0.36 | 1 | AXXATGG | 1651 | 2241 | 197 | Yes | MEMKQAMMMVESGSAAKLPTIVSSVSMQNIQPRGPARRRAIPSISKSREE  EIRKILRNNLQKTRQRLRSYSRHDLMIDPFEDNVSEIRFRKQRVEMERRM  SHYLTVPANRQETPPVRKVCFEPEHQVYTYDDSESTRGPTAQPNLSPADA  VSLLNESPQRPSQRGDGELRAKDEQEELKLSRCLSDPGPNKEEEGDG | 22.6 |
| 7 | 0.34 | 1 | AXXATGG | 475 | 2241 | 589 | Yes | MGTILVFAVVGTLWNAFFIGGMMYGVCQIEGAKLANVDLLSCLLFGSIIS  AVDPVAVLAVFEEIHINELLHILVFGESLLNDAVTVVLYHLFKEFSQAGT  VTVVDAVLGVVCFFVVSLGGVMVGAIYGLLGAFTSRFTSHTRVIEPLFVF  LYSYMAYLSAEVFHLSGIMSLIACGVMMRPYVEANISHKSYTTIKYFLKM  WSSVSETLIFIFLGVSTVAGPHAWNWTFVVSTVVLCLVSRVLGVIGLTFI  INKFRIVKLTKKDQFIVAYGGLRGAIAFSLGFLLTNNEMKHLFLTAIITV  IFFTVFVQGMTIRPLVELLAVKRKKENKGSINEEIHTQFLDHLLTGIECI  CGHYGHHHWKDKLNRFNKAYVKKWLIAGERSTEPQLISFYNKMEMKQAMM  MVESGSAAKLPTIVSSVSMQNIQPRGPARRRAIPSISKSREEEIRKILRN  NLQKTRQRLRSYSRHDLMIDPFEDNVSEIRFRKQRVEMERRMSHYLTVPA  NRQETPPVRKVCFEPEHQVYTYDDSESTRGPTAQPNLSPADAVSLLNESP  QRPSQRGDGELRAKDEQEELKLSRCLSDPGPNKEEEGDG | 66.3 |
| 15 | 0.30 | 1 | GXXATGa | 1003 | 2241 | 413 | Yes | MMRPYVEANISHKSYTTIKYFLKMWSSVSETLIFIFLGVSTVAGPHAWNW  TFVVSTVVLCLVSRVLGVIGLTFIINKFRIVKLTKKDQFIVAYGGLRGAI  AFSLGFLLTNNEMKHLFLTAIITVIFFTVFVQGMTIRPLVELLAVKRKKE  NKGSINEEIHTQFLDHLLTGIECICGHYGHHHWKDKLNRFNKAYVKKWLI  AGERSTEPQLISFYNKMEMKQAMMMVESGSAAKLPTIVSSVSMQNIQPRG  PARRRAIPSISKSREEEIRKILRNNLQKTRQRLRSYSRHDLMIDPFEDNV  SEIRFRKQRVEMERRMSHYLTVPANRQETPPVRKVCFEPEHQVYTYDDSE  STRGPTAQPNLSPADAVSLLNESPQRPSQRGDGELRAKDEQEELKLSRCL  SDPGPNKEEEGDG | 47.4 |

### Supplementary Figures

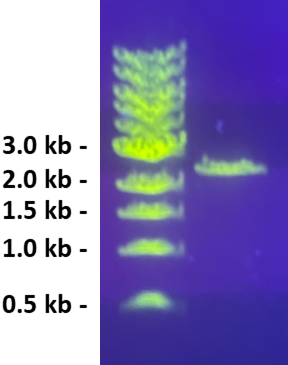

**Figure S1. The white seabass red blood cell (RBC) Slc9a1b PCR product (2372 bp).** Full-length cDNA sequences were obtained in 35 cycles of PCR reactions with Phusion DNA polymerase and specific primers (F: 5’TCC CGT ACT ATC CTC ATC TTC A-3’ R: 5’-CCT CTG CTC TCT GAA CTG TAA AT-3’) designed against the sequence of the phylogenetically characterized white seabass β-NHE (Fig. 3) obtained from the combined gill and RBC transcriptome. Gel electrophoresis (agarose 0.5% w/v) was for one hour at 4.5 V cm^-1^ in TAE buffer (Tris-acetate-EDTA) containing 1:10,000 SYBR-Safe (Thermo Fisher S33102; Waltham, USA) and the image was taken under short-wavelength ultraviolet light.

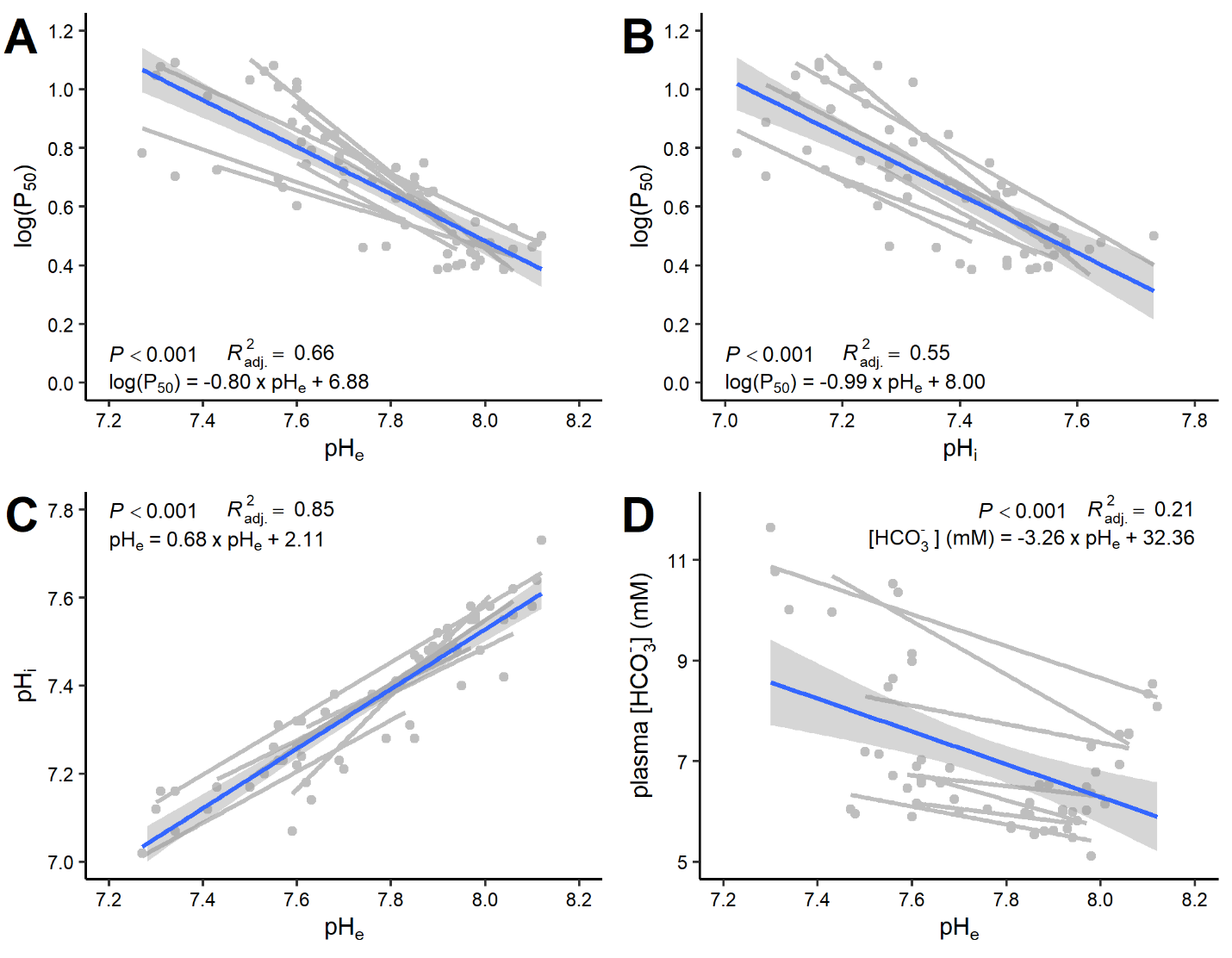

**Figure S2 Blood parameters determined by regression analysis in white seabass whole blood during hypercapnic acidification (PCO_2_ 0.3-2.5** **kPa).** A) the Bohr coefficient for extracellular pH (pH_e_) was calculated as the average slope of Δlog(P_50_) over ΔpH_e_ (P_50_ is the partial pressure of O_2_ that yields 50% Hb-O_2_ saturation); B) the Bohr coefficient for intracellular pH (pH_i_); C) the relationship between pH_i_ and pH_e_; and C) the non-bicarbonate buffer capacity of the blood was calculated as the average slope of plasma [HCO_3_^-^] over pH_e_. Grey data points represent individual measurements and a regression was plotted for each individual fish (*N* = 8). The blue regression line was fitted to the pooled dataset and the shaded area represents the 95% confidence intervals. The average slopes and s.e.m. were used to calculate the blood parameters shown in Figure 1.

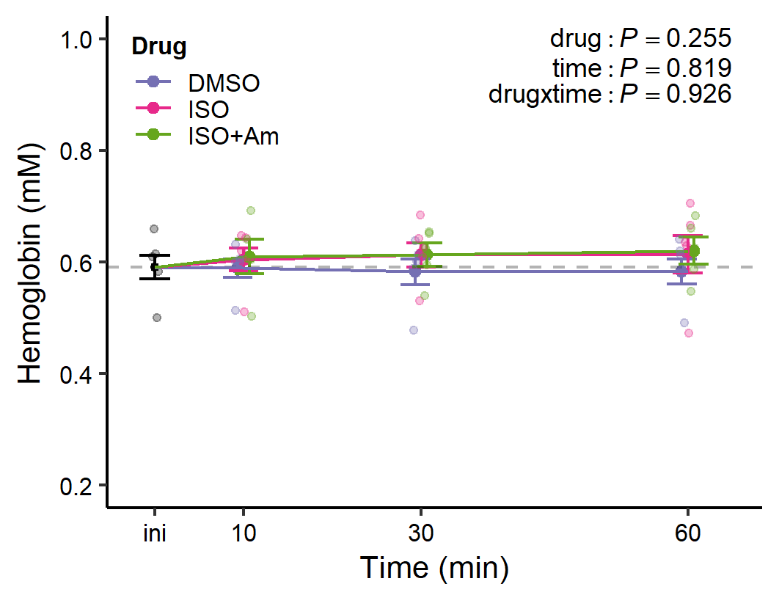

**Figure S3 No changes in hemoglobin concentration ([Hb]; mM) after adrenergic stimulation of white seabass whole blood.** Blood was equilibrated in tonometers at 3 kPa PO_2_ and 1 kPa PCO_2_ and treated with either: i) a carrier control (DMSO; 0.25%), ii) the β-adrenergic agonist isoproterenol (ISO; 10 µM final concentration) or iii) ISO plus amiloride (ISO+Am; 1 mM), an inhibitor of sodium-proton exchangers (NHE). The main effects of treatment (treat), time and their interaction term (inter) were analysed with a two-way ANOVA (*P* < 0.05, *N* = 6 and *N* = 5 for ISO+Am). There were no significant changes in [Hb] throughout the trials and therefore, the average concentration was reported in Figure 2 panel B. All data are means±s.e.m..

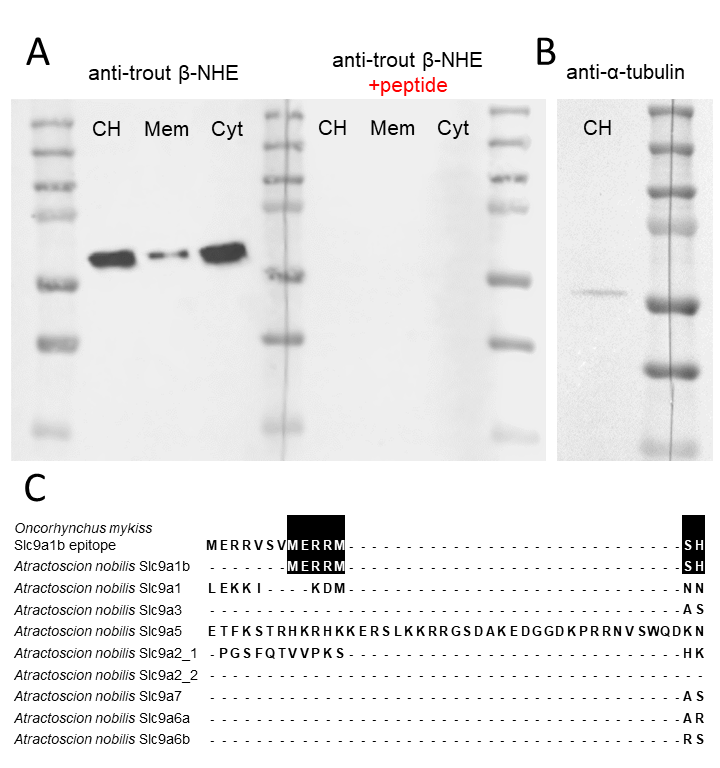
**Figure S4 Validation of the β-NHE antibody used for red blood cell (RBC) immunocytochemistry.** A) In Western blots, lanes of a gel were loaded with 5 µg protein from RBC crude homogenate (CH), membrane-enriched fraction (Mem) and cytosolic fraction (Cyt). Primary incubations were with 0.42 ng ml^-1^ of the polyclonal rabbit anti-trout β-NHE antibody and imaging was for a long exposure of 200 s. The β-NHE antibody labelled a single band with a predicted size of 66 kDa and longer exposures or higher protein loading did not reveal any additional bands. In peptide pre-absorption controls, the lanes were loaded with the same amount of protein, but were probed with primary antibody that was incubated over-night with its immune peptide, fully eliminating immuno-reactivity. B) A single lane was loaded with 60 µg RBC CH protein and was incubated with 4.7 ng ml^-1^ of the monoclonal anti-*Tetrahymena* α-tubulin antibody that recognized a single band of 54 kDa. C) The epitope sequence of the polyclonal anti-rainbow trout β-NHE antibody aligned with the nine white seabass NHE isoforms detected in the combined gill and red blood cell transcriptome and identified by phylogenetic analysis (Fig. 3). Homologous amino acid residues are highlighted in black and the corresponding region of the white seabass β-NHE sequence shared 50% sequence identity with the antibody epitope (7/14 consecutive amino acids). None of the other eight NHE isoforms in white seabass shared any amino acids within the epitope region, thus supporting the specificity of the antibody for the detection of white seabass β-NHE protein only.

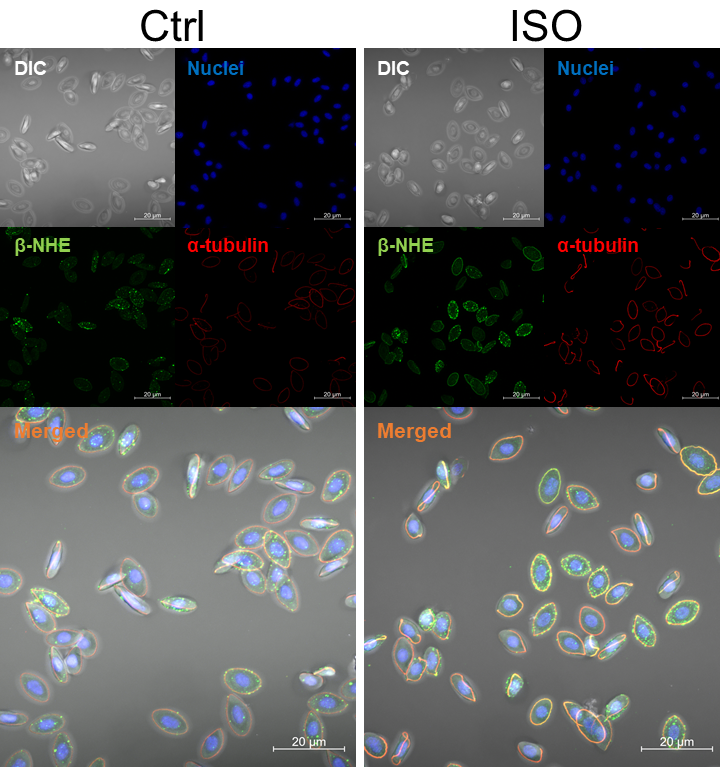

**Figure S5 Immunocytochemical localization of the β-adrenergic sodium-proton exchanger (β-NHE) in white seabass red blood cells (RBC).** Blood was equilibrated in tonometers at 3 kPa PO_2_ and 1 kPa PCO_2_ for 60 mins in the presence of the carrier control (DMSO; 0.25%), or the β-adrenergic agonist isoproterenol (ISO; 10 µM final concentration). Fixed cells were stained with a monoclonal anti-*Tetrahymena* α-tubulin antibody to visualize the marginal band (red), a polyclonal anti-trout β-NHE antibody (green), DAPI to visualize the cell nuclei (blue). Confocal immunofluorescence images were overlayed with differential interference contrast (DIC) showing cell morphology. The merged channels images revealed mostly intracellular signal for β-NHE in Ctrl RBCs, whereas in ISO-treated RBCs β-NHE protein co-localized (orange) with α-tubulin at the cell membrane.

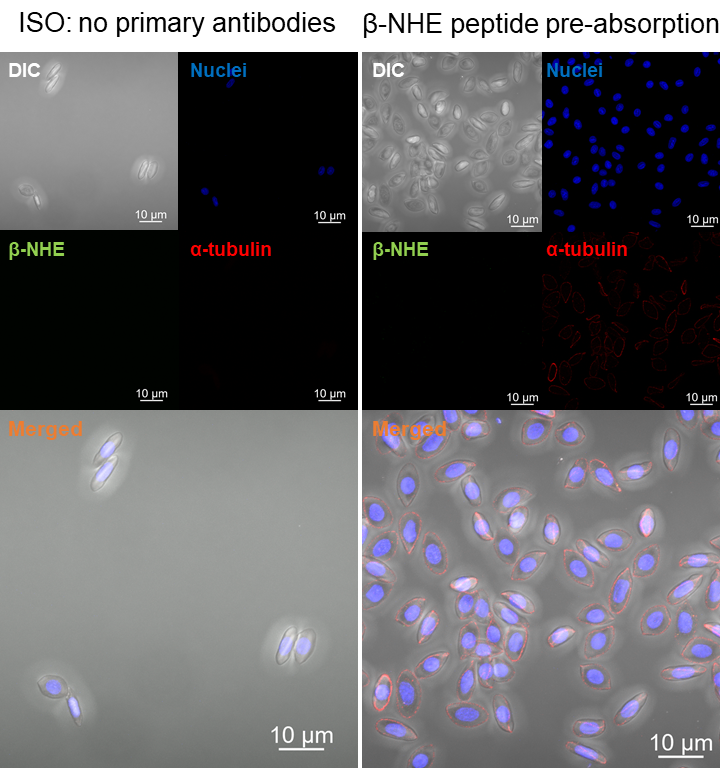

**Figure S6 Antibody controls for the immunocytochemical localization of proteins in white seabass red blood cells.** Blood was equilibrated in tonometers at 3 kPa PO_2_ and 1 kPa PCO_2_ for 60 mins in the presence of the β-adrenergic agonist isoproterenol (ISO; 10 µM). Fixed cells were incubated without primary antibodies or with a polyclonal anti-trout β-NHE antibody that was incubated with an excess of pre-immune peptide (1:10) overnight. Secondary antibodies were used according to standard protocols and DAPI was used to visualize the cell nuclei (blue). Confocal immunofluorescence images were overlayed with differential interference contrast (DIC) showing cell morphology. In the no-primary controls, the merged channels image validated the absence of immunoreactivity for β-NHE (green) and α-tubulin proteins (red), whereas the peptide pre-absorption control validated the absence of immunoreactivity for β-NHE (green), while α-tubulin, nuclei and cell morphology were clearly visible.

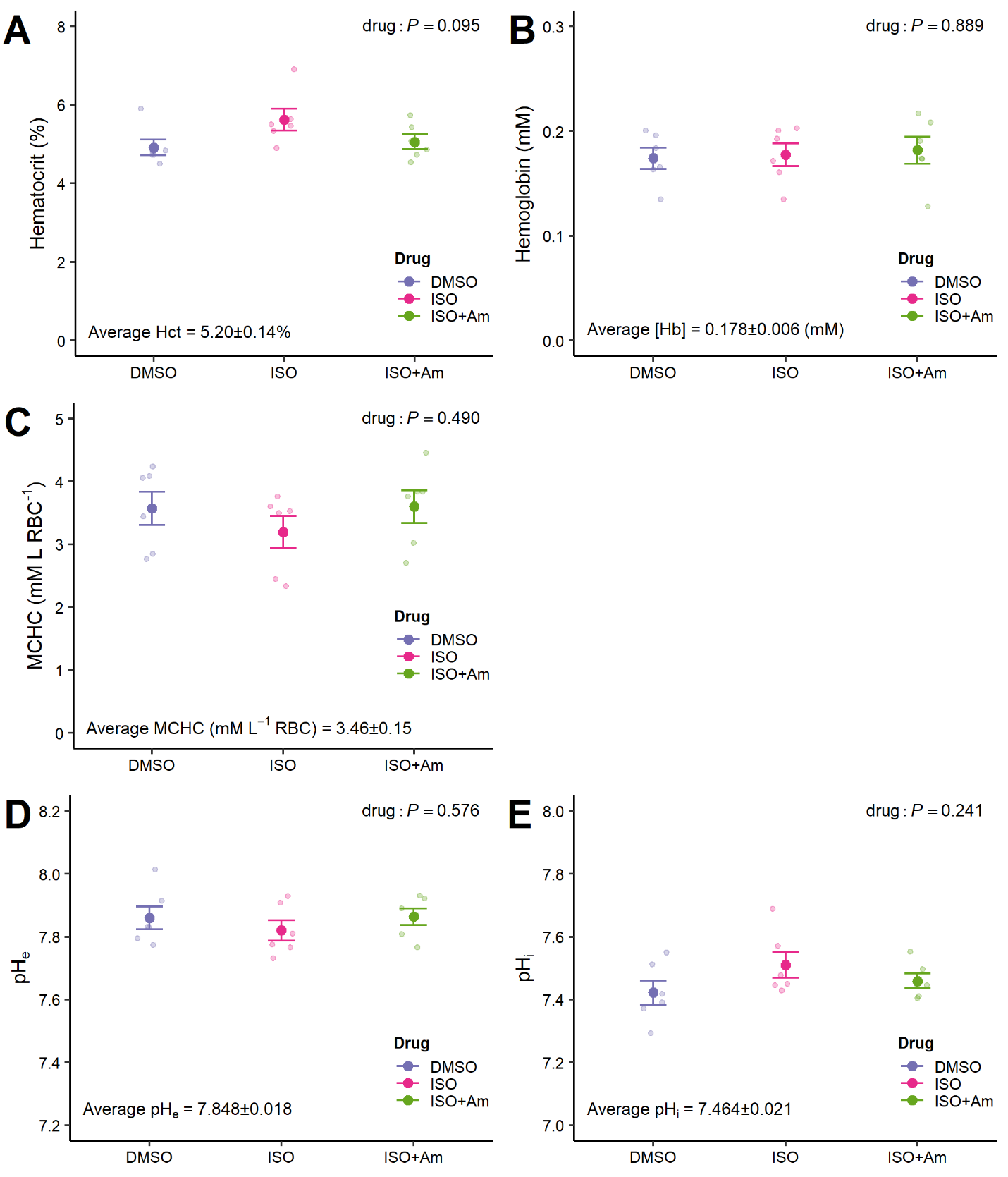

**Figure S7 Initial blood parameters before the spectrophotometric measurement of changes in Hb-O_2_ saturation during a respiratory acidosis.** A) Hematocrit (%), B) hemoglobin concentration (mM), C) mean cell hemoglobin content (MCHC; mM hemoglobin l^-1^ red blood cells), D) extracellular pH (pH_e_) and E) red blood cell intracellular pH (pH_i_). Blood was prepared in plasma at a Hct of 5%, equilibrated in tonometers at 21 kPa PO_2_ and 0.3 kPa PCO_2_ and treated with either: i) a carrier control (DMSO; 0.25%), ii) the β-adrenergic agonist isoproterenol (ISO; 10 µM), or iii) ISO plus amiloride (ISO+Am; 1 mM), an inhibitor of sodium-proton exchangers (NHE). The main effect of drug treatments (drug) was analysed with a One-way ANOVA (*P* < 0.05, *N* = 6). As there were no significant effects of drug treatments on these blood parameters, the average values during the trials are shown at the bottom of each panel. All data are means±s.e.m..

### Supplementary Movies

**Movie S1 Three-dimensional (3D) reconstruction of a control white seabass red blood cell immunostained for the β-adrenergic sodium proton exchanger (β-NHE).** Blood was equilibrated in tonometers at 3 kPa PO_2_ and 1 kPa PCO_2_ for 60 mins in the presence of the carrier control (DMSO; 0.25%). Fixed cells were stained with a monoclonal anti-*Tetrahymena* α-tubulin antibody to visualize the marginal band (red), a polyclonal anti-trout β-NHE antibody (green) and DAPI to visualize the cell nuclei (blue); co-localization of α-tubulin and β-NHE proteins is shown in orange. Super-resolution images were generated with the Zeiss AiryScan detector system and optical sectioning and 3D rendering was with the Imaris software. Intense β-NHE signal was detected in intracellular vesicle-like structures, with weaker membrane staining associated with α-tubulin in the RBC’s marginal band.

**Movie S2 Three-dimensional (3D) reconstruction of an adrenergically-stimulated white seabass red blood cell immunostained for the β-adrenergic sodium proton exchanger (β-NHE).** Blood was equilibrated in tonometers at 3 kPa PO_2_ and 1 kPa PCO_2_ for 60 mins in the presence of the β-adrenergic agonist isoproterenol (ISO; 10 µM final concentration). Fixed cells were stained with a monoclonal anti-*Tetrahymena* α-tubulin antibody to visualize the marginal band (red), a polyclonal anti-trout β-NHE antibody (green) and DAPI to visualize the cell nuclei (blue); co-localisation of α-tubulin and β-NHE proteins is shown in orange. Super-resolution images were generated with the Zeiss AiryScan detector system and optical sectioning and 3D rendering was with the Imaris software. Intense β-NHE signal was associated with α-tubulin in the RBC’s marginal band with only weak intracellular staining.
